# Structural insights into peptidoglycan hydrolysis by the FtsEX system in *Escherichia coli* during cell division

**DOI:** 10.1101/2024.01.16.575974

**Authors:** Jianwei Li, Yutong He, Xin Xu, Martin Alcorlo, Jian Shi, Souvik Naskar, Nicholas S. Briggs, David I. Roper, Juan A. Hermoso, Lok-To Sham, Min Luo

## Abstract

Bacterial cell division relies on precise peptidoglycan (PG) remodelling, a process orchestrated by the FtsEX complex. Comprised of FtsE and FtsX, this complex collaborates with EnvC, a periplasmic lytic enzyme activator, to regulate septal PG hydrolysis by amidases like AmiB. While recent structural investigations, particularly of *Pseudomonas aeruginosa* FtsEX (*Pae*FtsEX), have shed light on complex interactions and proposed activation mechanisms, the structural intricacies governing PG degradation by the FtsEX complex and EnvC in *Escherichia coli* cytokinesis remain unexplored. In this study, we present a comprehensive biochemical and structural analysis of *E. coli* FtsEX complexes, unveiling a key role for ATP in complex stabilization that extends across bacterial species. Upon EnvC binding, ATPase activity markedly increases. High-resolution structures of *Eco*FtsEX, both in the presence and absence of EnvC, reveal a symmetrical conformation of *Eco*FtsEX capable of accommodating the inherent asymmetry of EnvC, mediated by flexible loops within the periplasmic domain. Our negative-staining imaging showcases an elongated *Eco*FtsEX/EnvC/AmiB complex reminiscent of the *Pae*FtsEX system. These findings collectively provide intricate insights into the regulation of PG cleavage by FtsEX in *E. coli* - a pivotal model system used in pilot genetic studies, suggesting a conserved mechanism for precise hydrolase activation in bacteria.

## Introduction

In most bacteria, peptidoglycan (PG) remodeling during cell division is a precisely regulated process that ensures daughter cells separate without damaging the cell wall (1–3). This process involves a set of essential proteins that coordinate PG synthesis and degradation. In Escherichia coli, at least ten proteins contribute to cell division, many of which are organized by the cytoskeletal protein FtsZ (4, 5). FtsZ recruits FtsA and FtsEX to form the early divisome, a protein assembly at the division site. After a short delay, additional proteins, including FtsK, FtsQLB, FtsWI, and FtsN, are recruited to the septum (4, 6). The arrival of FtsN is a critical step that initiates cell constriction, likely through activating the FtsQLB complex and FtsA (4, 7–9). The final step in division, daughter cell separation, is carried out by PG-cleaving enzymes called amidases (AmiA, AmiB, and AmiC). These enzymes require activation by coiled-coil domain proteins: EnvC (which activates AmiA and AmiB) and NlpD (which activates AmiC) (1, 3, 10, 11). If these systems are simultaneously inactivated, the daughter cells will be unable to separate and form long chains (12).

PG remodelling is linked to cytokinesis by an ATP-binding cassette (ABC) transporter-like protein complex FtsEX. Rather than transporting a substrate, the FtsEX complex transduces conformational changes that span the cytoplasmic membrane (13, 14). Driven by ATP hydrolysis of FtsE, these changes are required for stimulating EnvC/AmiA/AmiB activities (15, 16) and initiating cell constriction via FtsA (15, 17–19). The FtsEX complex also recruits other cell division proteins to the divisome (17, 19). Thus, deletions of *ftsEX* result in non-viable cells unless they are grown in permissive conditions, such as at a low temperature or in media with high osmolarity (20, 21). Alternatively, the essentiality of *ftsEX* can be bypassed by gain-of-function mutations that strengthen the interaction between FtsN and FtsA (17, 20).

Several partial structures of the FtsEX complex have been reported, some are with their interacting partners (22–25). Very recently, progress has been made in elucidating the full-length structure of the FtsEX complex from *Pseudomonas aeruginosa* (*Pae*FtsEX), both in the presence and absence of critical regulators like ATP, EnvC, and AmiB (26). These structures offer insight into the interfaces between FtsE, FtsX, and their binding partners. In addition, they suggest a mechanism in which the rearrangement of regulatory helices of EnvC and AmiB activates peptidoglycan hydrolysis at the septum (26). Yet, the majority of work regarding FtsEX was done in *Escherichia coli* (15–21), where an abundance of genetic studies has laid a significant foundation for our understanding of PG degradation activation in bacterial cell division. However, the intricate molecular basis orchestrating this process remains elusive, underscoring the need for a direct structural and biochemical investigation within this extensively studied model system. Specifically, given the relatively low sequence similarity between the FtsEX complexes from these two species, it is unclear whether the structural information of *Pae*FtsEX can reconcile the findings from genetic studies. Furthermore, previous genetic studies have proposed a direct interaction between FtsEX and components of the constriction ring, such as FtsA and FtsZ (18, 27), to ensure the accurate localization of FtsEX at the division site. However, there is currently a lack of supporting biochemical and structural evidence in this regard.

In this study, we present a comprehensive analysis of the structure and function of the *Escherichia coli* FtsEX complex (*Eco*FtsEX) in conjunction with EnvC, focusing on its regulation of AmiB activation for PG cleavage. Our investigation uncovers a previously unrecognized function of ATP in stabilizing the FtsEX complex, a role that appears to be conserved across different bacterial species. Notably, we observe a substantial increase in the ATPase activity of *Eco*FtsEX upon binding with EnvC. High-resolution structures of *Eco*FtsEX alone and with EnvC were obtained at 3.9 and 3.4 Å, respectively, offering insights into the EnvC recognition mechanism. Interestingly, *Eco*FtsEX maintains a symmetrical conformation even when complexed with the asymmetrical EnvC. Comparative analyses identify the periplasmic domain’s flexible loops as key to EnvC interaction, revealing structural adaptability. Furthermore, our negative-staining imaging experiments unveil an elongated conformation of the *Eco*FtsEX/EnvC/AmiB complex for PG cleavage, reminiscent of the FtsEX system from *Pseudomonas aeruginosa*. Together, our results revealed a detailed regulation of PG cleavage by FtsEX in the key model system of *E. coli*, suggesting a conserved mechanism for hydrolase activation in bacteria.

## Results

### ATP Binding as a Universal Stabilizing Factor for the FtsEX Complex

In our efforts to reconstitute the full-length *Eco*FtsEX complex in vitro, we explored various strategies involving changing the expression constructs and extraction conditions. During the purification process, we observed that the protein remained stable in the presence of 10% glycerol. However, almost all protein precipitated out after the removal of glycerol either through size-exclusion chromatography (SEC) or during the peptidisc reconstitution (**Figure 1A**). This indicated a low level of stability for the complex. To investigate whether the stability of the FtsEX complex might be dependent on its cognate partner, EnvC, we conducted co-expression experiments by introducing EnvC along with FtsE and FtsX (**Figure 1A**). Despite the presence of EnvC, FtsEX still exhibited a tendency to precipitate.

**Figure 1.**
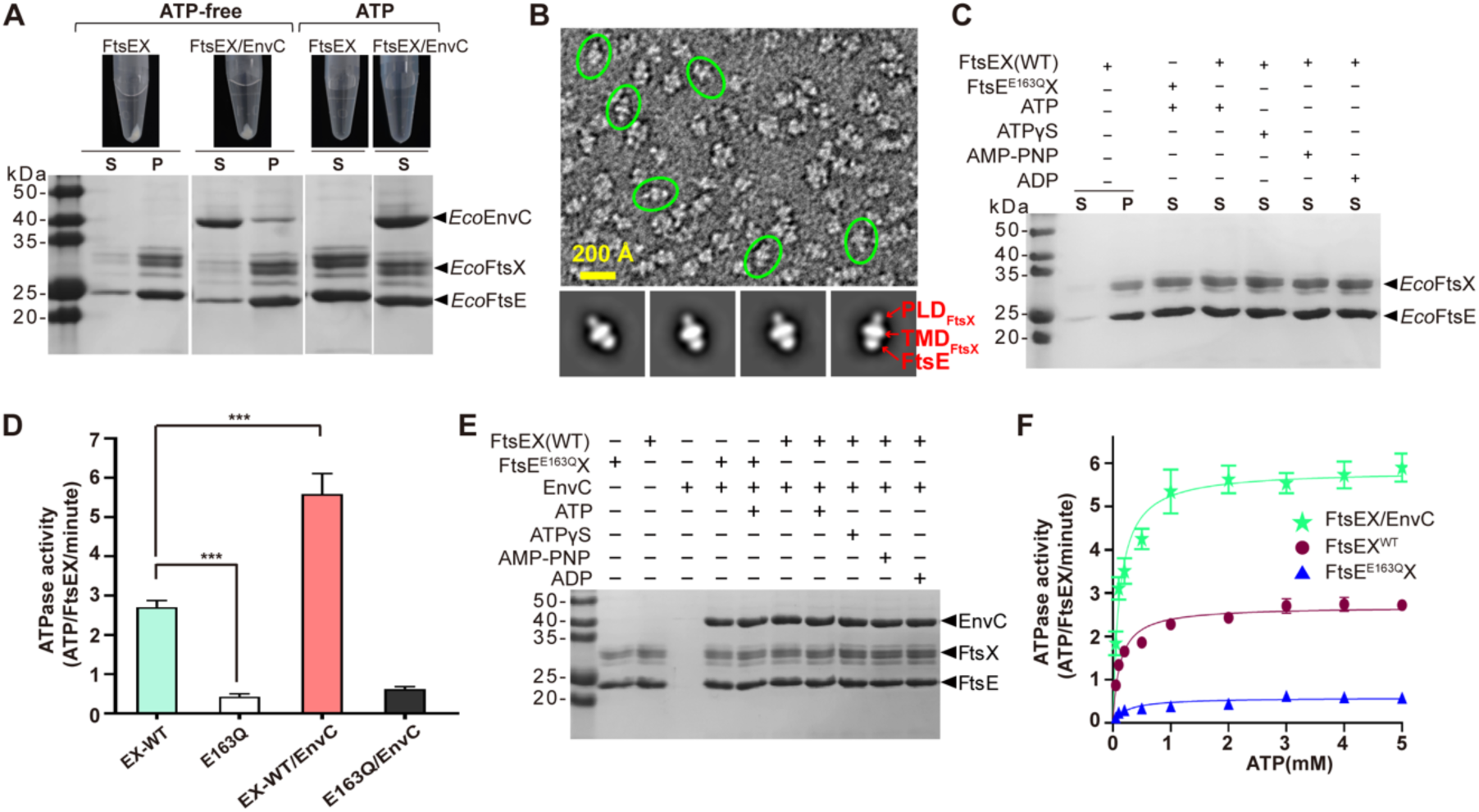
Biochemical reconstitution of *Eco*FtsEX and *Eco*FtsEX/EnvC complexes. **(A)** Evaluation of ATP’s impact on FtsEX and FtsEX/EnvC complexes. Purified FtsEX and FtsEX/EnvC complexes undergo precipitation in the absence of ATP but form stable complexes when ATP is present during purification. S: Supernatant, P: Precipitate. **(B)** Representative negative staining image and 2D average of FtsEX samples purified in the presence of ATP, with FtsEX highlighted in green. The FtsE subunit, Transmembrane Domain (TMD), and Periplasmic Domain (PLD) of FtsX are clearly visible. **(C)** Assessment of ATP and ATP analogs’ influence on the stability of FtsEX complex formation. ATP and ATP analogs contribute to the stabilization of the FtsEX complex. S: Supernatant, P: Precipitate. (**D**) Assessment of ATPase activity in FtsEX and FtsEX mutants, with and without EnvC. Notably, the FtsE^E163Q^X mutants exhibit significantly reduced ATPase activity. Data represent an average of three replicates, with error bars indicating mean ± SD. Statistical analysis performed using a two-tailed unpaired t-test; ***P < 0.0005. (**E**). Pull-down study illustrating EnvC binding to FtsEX or its ATPase mutant in the presence or absence of ATP or ATP analogs. The initial FtsEX or FtsE^E163Q^X added to the resin contains 2 mM ATP, the ATP was subsequently removed by the wash buffer. (**F**) ATPase activity comparison between FtsEX and the FtsEX/EnvC complex in peptidiscs. Data represent an average of three replicates, with error bars indicating mean ± SD.

Given the persistent challenges in achieving a stable FtsEX complex, we redirected our focus towards exploring the potential role of the substrate ligand, ATP, in complex stabilization. ATP, well-known for its role as a biological energy source, has also been associated with the stabilization of proteins, as reported in recent studies (28, 29). To investigate this hypothesis, we conducted a series of experiments to assess the impact of ATP on the FtsEX or the FtsEX/EnvC complex (**Figure 1A**). Remarkably, the introduction of ATP resulted in the formation of a robust *Eco*FtsEX complex, both in its standalone form and in the presence of EnvC. This complex remained tightly associated throughout the SEC process (**Figure supplement 1A**). Further validation through negative-staining electron microscopy confirmed that the purified *Eco*FtsEX complex maintained stability, with clearly discernible features corresponding to an intact ABC transporter containing two FtsE and two FtsX subunits (**Figure 1B**). This compelling evidence suggests that ATP, acting as the substrate of FtsEX, also plays a pivotal role in stabilizing this crucial complex formation.

Interestingly, ATP analogues, including ATPγS, AMP-PNP, and ADP, also exhibited similar stabilizing effects (**Figure 1C**). As they are not hydrolysable for FtsEX, it indicates that ATP binding, rather than hydrolysis, is sufficient for complex stabilization. This was corroborated by the introduction of a FtsE mutant (E163Q), inspired by related ABC transporters (26), which is incapable of ATP hydrolysis. When this mutant was assessed for complex formation with FtsX in the presence of ATP, it indeed formed a stable complex (**Figure 1C**). These findings suggest ATP binding alone is responsible for stabilizing the complex formation.

To expand the scope of our investigation, we explored whether ATP could similarly stabilize FtsEX complexes from other bacterial species, which had also presented challenges in in-vitro reconstitution. We expressed and purified FtsEX from various species, including *Streptococcus pneumoniae, Pseudomonas aeruginosa*, and *Bacillus subtilis,* in the presence and absence of ATP. Remarkably, regardless of the bacterial species, ATP consistently enhanced the stability of the complexes, as demonstrated by either increased complex yield in SEC elution profiles or with a more accurate stoichiometry between FtsE and FtsX as shown from the SDS-page gel (**Figure supplement 1B**). Collectively, our findings underscore the pivotal role of ATP in stabilizing FtsEX complexes. Notably, the ability of other nonhydrolizable ATP analogues to replicate this effect suggests that ATP binding is a crucial step required to facilitate the coupling of FtsX and FtsE for the formation of a functionally stable complex.

### Biochemical Characterization of FtsEX and Its Complexes with EnvC in *E. coli*

The His-tagged full-length FtsE and FtsX proteins from *E. coli* (str. K-12, MG1655) were overexpressed in *E. coli* BL21(DE3) and subsequently purified using a combination of affinity and size-exclusion chromatography (SEC) (**Figure supplement 1A**). For subsequent functional and structural investigations, FtsEX was solubilized in DDM and reconstituted into peptidiscs (see Methods). Given the proven stabilization role, 0.5 mM ATP was maintained throughout the entire purification process of *Eco*FtsEX.

Assays assessing ATPase activity demonstrated that, in contrast to previously reported FtsEX complexes from *Pseudomonas aeruginosa*, the *Eco*FtsEX complex exhibited obvious basal ATPase activity (**Figure 1D**). To confirm this, we introduced the FtsE mutant (E163Q) as a control, which was not able to hydrolyse ATP. As anticipated, ATPase activity in the mutant was markedly diminished (**Figure 1D**), while low ATPase activity was still detected, potentially attributable to co-purified proteins.

To understand the role of ATP binding/hydrolysis on EnvC recruitment, a protein pull-down assay was conducted. As ATP was present in the purified protein, a wash step with buffer in the presence of Mg^2+^ and 10% glycerol was performed to remove the extra ATP for the FtsEX complex that bind to the resin. Next, bound FtsEX and EnvC were mixed in the presence of ATP or its non-hydrolysable analogue, ATPγS (**Figure 1E**). The results revealed that EnvC stably associated with FtsEX, with its binding seemingly unaffected by ATP hydrolysis. This was evidenced by the comparable amounts of EnvC bound in the presence of 3 mM ATP, as well as its non-hydrolysable analogs ATPγS and AMP-PNP, or even the hydrolysed product ADP. To confirm this, FtsEX^E163Q^ mutant was introduced. Results, similar to those from the wild type, showed that the mutant still demonstrated similar EnvC binding levels. The comparable binding of EnvC to FtsEX in the presence of ATP, non-hydrolyzable analogs (ATPγS, AMP-PNP), and ADP suggests that ATP binding is not required for EnvC recruitment to FtsEX, which aligns with multiple previous studies that show the PLD domain of FtsEX is able to interact with EnvC in the absence of FtsE (30). This indicates that EnvC association with FtsEX is nucleotide-independent and does not rely on ATP hydrolysis. Instead, ATP binding likely plays a role in stabilizing the FtsEX complex, rather than directly regulating EnvC recruitment.

Additionally, the effects of EnvC binding on the ATPase activity of *Eco*FtsEX were probed (**Figure 1D**). An enhanced activity was observed upon the addition of EnvC. This EnvC-mediated increase in ATPase activity was further validated through ATPase assays involving varying ATP concentrations for both FtsEX and the FtsEX/EnvC complex. Both sets of data fit well with the Michaelis-Menten equation (**Figure 1F**). Notably, in the presence of EnvC, the V_max_ (92.7±5.1 nmol/mg/min) was approximately 2.2-fold higher than that observed for FtsEX (42.7±2.1 nmol/mg/min) alone. However, no large differences were observed regarding the K_m_ (0.11±0.02 mM for FtsEX/EnvC; 0.13±0.02 mM for FtsEX). Together, these findings underscore that in the presence of ATP, FtsEX and EnvC form a robust complex; while *Eco*FtsEX inherently exhibits ATPase activity, the presence of EnvC stimulates this activity.

### CryoEM structure of *Eco*FtsEX reveals a symmetrical PLD arrangement

To elucidate the full-length structure of *Eco*FtsEX, we subjected the WT *Eco*FtsEX sample, incubated with 3 mM ATPγS, to single-particle CryoEM analysis and achieved a resolution of 3.7 Å (**Figure supplement 2**). This map was of sufficient resolution for detailed model-building for the Transmembrane domains (TMD), the nucleotide-binding domains (NBD), and a large portion of the Periplasmic domain (PLD); in addition, density for two bound nucleotides was clearly captured as well (**Figure supplement 3A**).

*Eco*FtsEX, akin to most Type VII ABC transporters, possesses one coupling helice (CH) and a C terminal tail (C-tail) (**Figure 2A**). Cryo-EM structure reveals the assembly of the full *Eco*FtsEX system (**Figure 2B**). This comprises two cytosolic FtsE NBD domains and two FtsX, with TMD domains embedded in the membrane and PLD extending into the periplasmic space. Both TMD and NBD regions in the *Eco*FtsEX structure exhibit canonical folds, similar to other characterized Type VII ABC transporters such as MacB, LolCDE, and *Pae*FtsEX. The overall conformation of *Eco*FtsEX is quite similar to that of *Pae*FtsEX (**Figure supplement 3B**). Notably, the N-terminal region of FtsX (Residues 53-70) forms an elbow helix (EH) oriented parallel to the membrane (**Figure 2B**). Like its counterpart in *Pae*FtsEX, the extreme N-terminal region of FtsX (Residues 1-52) was not resolved, suggesting its highly disordered nature.

**Figure 2.**
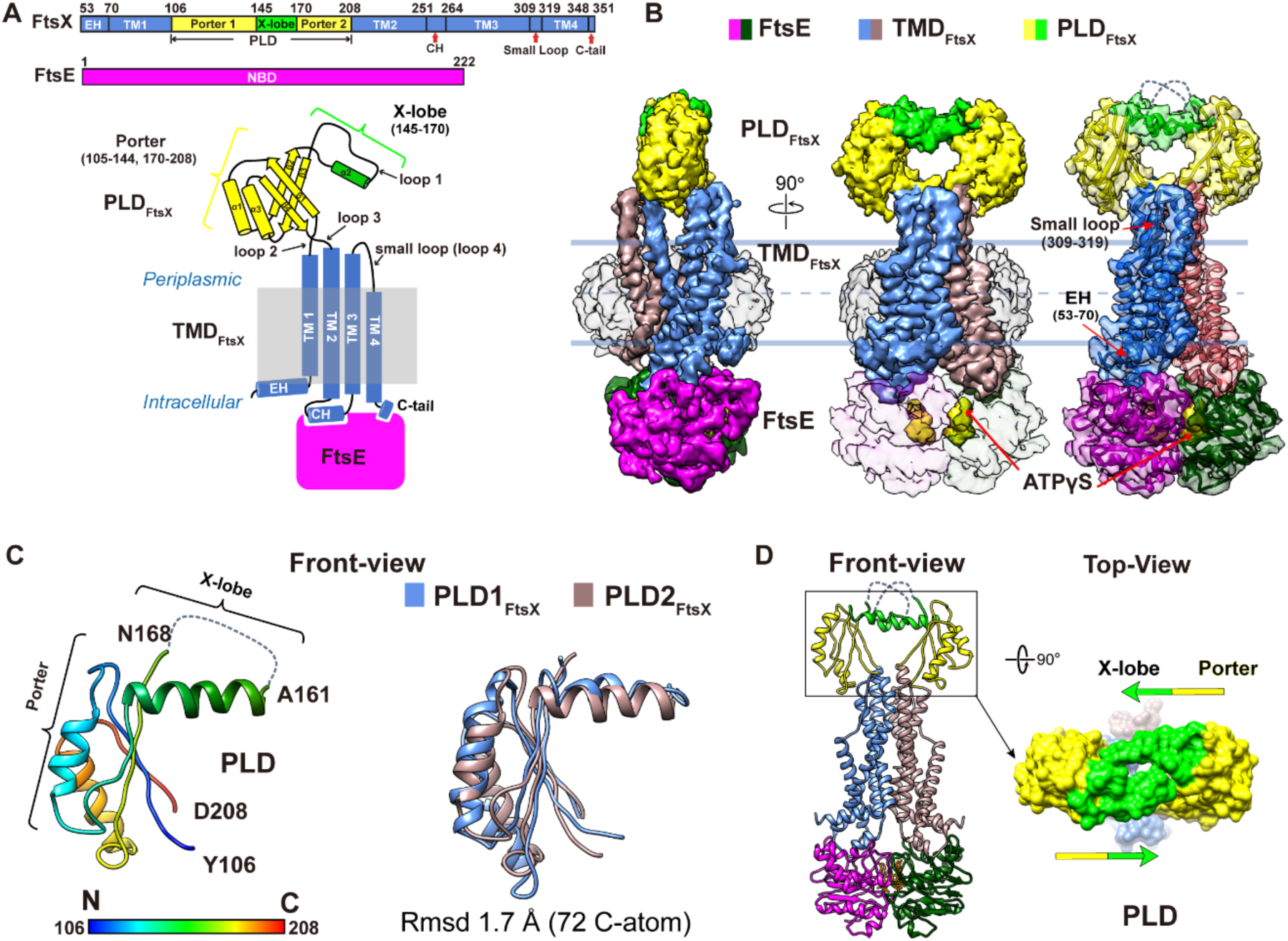
Cryo-EM study of *Eco*FtsEX complex in peptidiscs. (**A**) Topological diagram of FtsEX, highlighting the α helices represented as cylinders and β sheets depicted as arrows. Color scheme: FtsE (magenta and dark green), FtsX (cornflower blue), Porter of PLD_FtsX_ (yellow), and X-lobe of PLD_FtsX_ (green). CH: coupling helix, EH: elbow helix. (**B**) Left and middle panels depict the side and front views of the density map of FtsEX, with ATPγS displayed in the front view at 90% transparency. Ribbon representation of WT FtsEX in the presence of ATPγS with EM density. Color scheme: FtsE (magenta and dark green), FtsX (cornflower blue and rosy brown), Porter of PCD_FtsX_ (yellow), and X-lobe of PLD_FtsX_ (green). EH: elbow helix. (**C**) Left panel presents a rainbow representation of the PLD domain of FtsX, transitioning from blue (N-terminal) to red (C-terminal). Right panel shows a front view overlay of two PLD domains. (**D**) Surface representations of the PLD domains in a top-view configuration, with the Porter of the PLD colored in yellow and the X-lobe in green. In *Eco*FtsEX, the Porter and X-lobe are positioned facing each other.

The entire PLD domain comprises a smaller loop and a larger domain. For ease of reference, only the larger domain is termed the PLD (**Figure 2A**). The smaller loop encompasses 14 residues between TM3 and TM4. The principal PLD domain, spanning residues between TM1 and TM2 (106–208), displays a canonical topology, with a Porter domain and an X-lobe (14, 22). Compared to the TMD and NBD regions, PLD domains exhibit higher flexibility, evident from their diminished EM density at a contour level of 0.22 in Chimera. Lowering the contour to 0.1 in Chimera reveals clear density, enabling the visualization of PLD’s secondary structures (**Figure supplement 2F**), with an exception for the X-lobe segment with residues 161-168 (**Figure 2C**). Structurally, the individual PLD domains are superimposable (**Figure 2C**), and also align closely with those in *Pae*FtsEX (**Figure supplement 4**), despite their amino acid sequence identities with *E. coli* being only 29%.

The PLD domain of *Eco*FtsEX structure offers a snapshot with well-defined secondary structures, highlighting a 2-fold symmetrical conformation (**Figure 2D**). The X-lobes of both PLDs are oriented mutually inwards with sideways blocked and appear to facilitate EnvC binding strictly from the top of the FtsEX complex, mirroring findings in the *Pae*FtsEX system.

### CryoEM structure of *Eco*FtsEX/EnvC Complex elucidates EnvC binding and ATP-Dependent LytM Release

To elucidate the EnvC recruitment by FtsEX in *E. coli*, purified FtsEX^E163Q^ mutant was reconstituted in peptidiscs and incubated with EnvC in the presence of 2 mM ATP for cryo-EM analysis (**Figure supplement 5**). After 3D reconstruction, a major conformation stands out with resolution determined at 3.4 Å (**Figure supplement 5F**). The quality of the EM density allowed modelling of nearly all sidechains of FtsEX, the two bound nucleotides, and a portion of EnvC’s coiled-coil domain (CCD) (**Figure 3A & Figure supplement 6**). The density of EnvC’s remaining regions, including part of the CCD domain, the LytM domain and the regulatory helix, was absent, suggesting significant flexibility. The global configuration of the FtsEX/EnvC complex is similar to the previously described *Pae*FtsEX system (**Figure supplement 7**), where EnvC inserts from above between the two PLD domains along the FtsEX central axis.

**Figure 3.**
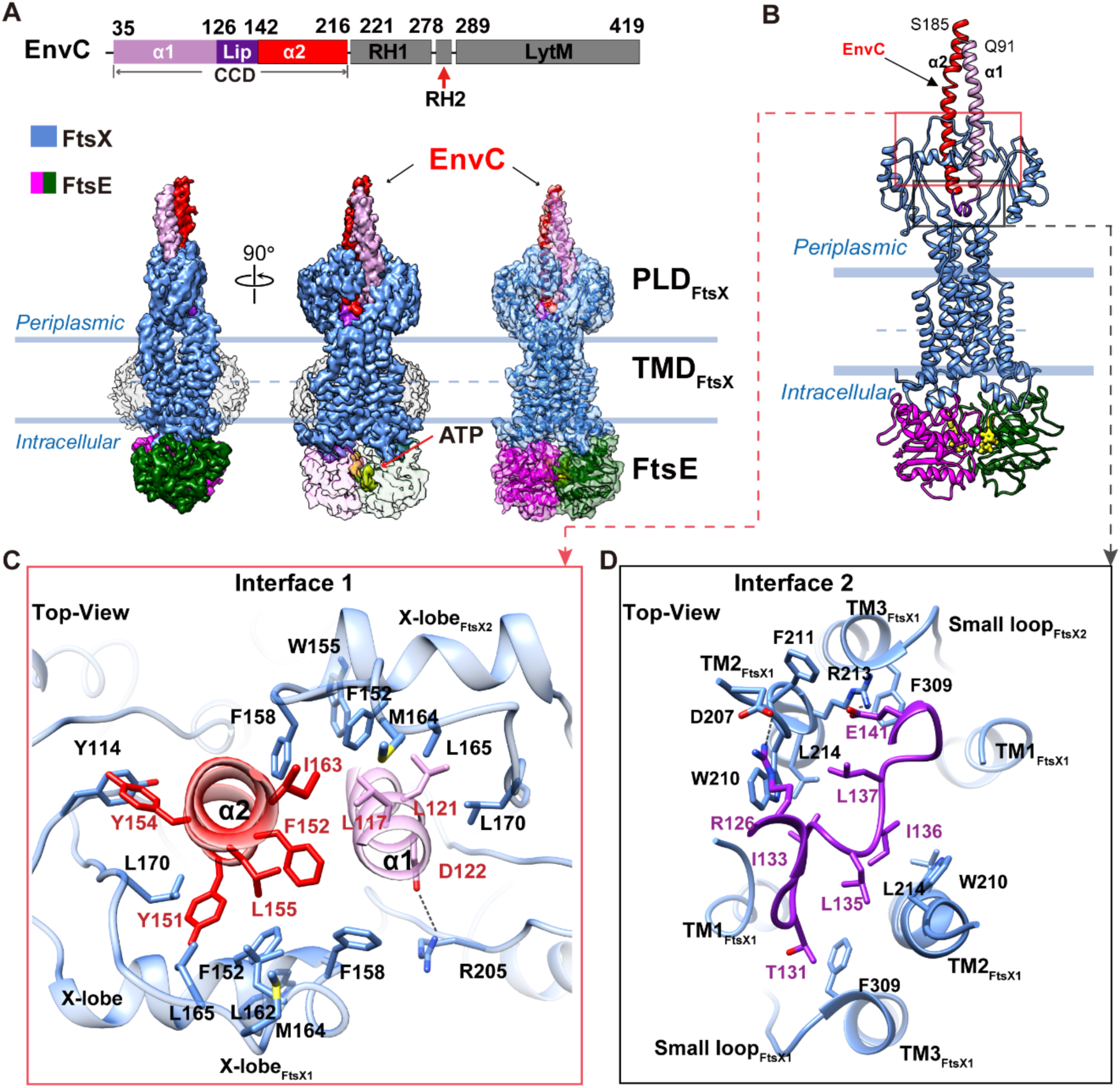
Cryo-EM structure of the EnvC-bound *Eco*FtsEX complex. (**A**) Domain arrangements and the overall structure of FtsEX/EnvC. Left and middle panels display side and front views of the cryo-EM density maps for FtsEX-EnvC complex in the presence of ATP. The right panel presents a ribbon representation of the structural model overlaid with the EM density. In the front-view, FtsE is depicted at 90% transparency to reveal the ATP-binding site. Color scheme of EnvC: α1 (plum), lip (purple), α2 (red), RH1, RH2, and LytM domains (grey). (**B**) EnvC binding to the PLD domain of FtsEX is depicted with ribbon representations. Color scheme: FtsX (cornflower blue), FtsE (magenta/dark green), and EnvC (red). (**C**) **&** (**D**). Top-down views provide a detailed depiction of the interaction between EnvC and FtsEX.

To dissect the state of the bound EnvC, we compared our cryoEM FtsEX/EnvC complex with the crystal structure of the PLD domain in complex with EnvC (PLD/EnvC) from *E. coli* (PDB code: 6TPI) (*22*), where EnvC was in an inactivated state. In the PLD/EnvC structure, a distinct asymmetrical conformation is evident: one PLD is positioned about 5 Å higher than the other one, accommodating the bound asymmetric EnvC with its LytM domain locked (**Figure supplement 8A, left**). However, in our full-length ATP-bound FtsEX/EnvC structure, the two PLDs are highly symmetrical (**Figure supplement 8A, right**). Besides this, the bound EnvC has also undergone conformational shifts (**Figure supplement 8B**). In contrast to the PLD/EnvC crystal structure, where the two CCD helices exhibit huge asymmetry (**Figure supplement 8B, left**), our structure reveals a more symmetrical arrangement induced by simultaneous compression from both PLD domains (**Figure supplement 8B, right**). Consequently, the distance between the terminal residue of the α1 helix (Q127) and the initial residue of the α2 helix (Q143) within the CCD domain is reduced from 24 Å to a mere 10 Å, while residing at a relatively similar height from the PLD domain. This compression of the CCD helices may drive the release of the LytM domain, supported by the lack of LytM density in our EM map. Together, we hypothesize that in the presence of ATP, the LytM domain is released, facilitating AmiB recruitment.

### Comparison Between *E. coli* and *P. aeruginosa* FtsEX/EnvC Structures

To further analyze the conservation and divergence of FtsEX/EnvC across species, we carefully compared our newly determined *E. coli* FtsEX/EnvC structure with the previously reported *P. aeruginosa* FtsEX/EnvC complex (**Figure supplement 9**). In both structures, the NBD domain (FtsE) dimerizes upon nucleotide binding, forming a similar overall dimeric assembly. This is coupled with a largely conserved conformation of the TMD domain (FtsX), with a root-mean-square deviation (RMSD) of 2.05 Å across 386 α-carbons of FtsE and 2.64 Å across 349 α-carbons of the FtsX TMD domain, indicating a high degree of structural similarity. Additionally, in both systems, EnvC inserts into the PLD domain in a vertical conformation, further supporting a conserved mode of interaction.

Despite these overall similarities, subtle differences exist between the two species. In *E. coli*, EnvC binds even more vertical relative to the FtsEX axis, with about 5° difference compared to the *Pae*FtsEX/EnvC structure. Along with this, a tighter coiled-coil (CC) conformation of *Eco*EnvC was observed, accompanied by a further compressed X-lobe in *E. coli*. The X-lobes, tightly coupled to the coiled-coil domains, appear to have been pushed further down toward the membrane in the *E. coli* structure. This may, in turn, induce a more compact CCD domain in EnvC, explaining the tighter conformation observed in *E. coli* compared to *P. aeruginosa*. These structural differences suggest potential species-specific variations in EnvC recruitment or activation. However, given the limited number of available FtsEX/EnvC structures, these conclusions remain preliminary. Additional structures from other species will be required to determine whether these variations reflect functional differences or simply structural plasticity within the FtsEX/EnvC system. Nonetheless, our study provides key insights into the core architecture and conserved interactions of FtsEX/EnvC, offering a structural framework for further investigations into bacterial cell division regulation.

### Detailed EnvC interactions with FtsEX

The ATP-bound *Eco*FtsEX/EnvC structure also revealed detailed interactions with EnvC. This interaction mainly involves two interfaces and covers an approximate surface area of 2290 Å^2^, predominantly consisting of hydrophobic contacts (**Figure 3B**).

The primary interface, also resolved in the PLD/EnvC crystal structure (22), features residues Y114, F152, W155, F158, L162, L165, and L170 of FtsX associating with residues L117, L121, L148, Y151, F152, Y154, L155, and I163 of EnvC (**Figure 3C**). Conversely, the secondary interface encompasses the Lip of CCD from EnvC, and the small loop of FtsX. Interacting residues at this interface comprise I133, L135, I136, L137, E141 from EnvC and W210, L214, F309 from FtsX. Although hydrophobic interactions dominate, several electrostatic interactions exist, notably the salt bridges (R126_EnvC_-D207_FtsX_, E141_EnvC_-R213_FtsX_) (**Figure 3D**). These interactions have largely been validated by previous studies (22), mutations like Y114E, F152E, and L165A in FtsX’s periplasmic domain notably affected EnvC’s binding efficacy, affirming their role in recruiting EnvC.

### Recognition of Asymmetrical EnvC by Symmetrical FtsEX: Unveiling Key Loops for Conformational Adaptivity

Intriguingly, our analysis of the FtsEX/EnvC structure showed that even when bound to the asymmetrical EnvC, FtsEX—including its two PLD domains interacting directly with EnvC— maintains a symmetrical conformation (**Figure 4A**). Compared to the *Eco*FtsEX structure without EnvC, which is also symmetrical, the only noticeable change is the outward shift of the two PLD hooks by about 12 Å (**Figure 4A**), a result of their interaction with the CCD domain of EnvC (**Figure 4B**). This observation poses an interesting question: How does the inherently symmetrical FtsEX cater to an asymmetrical binding partner? To explore this, we closely analysed the FtsX-EnvC binding interface.

**Figure 4.**
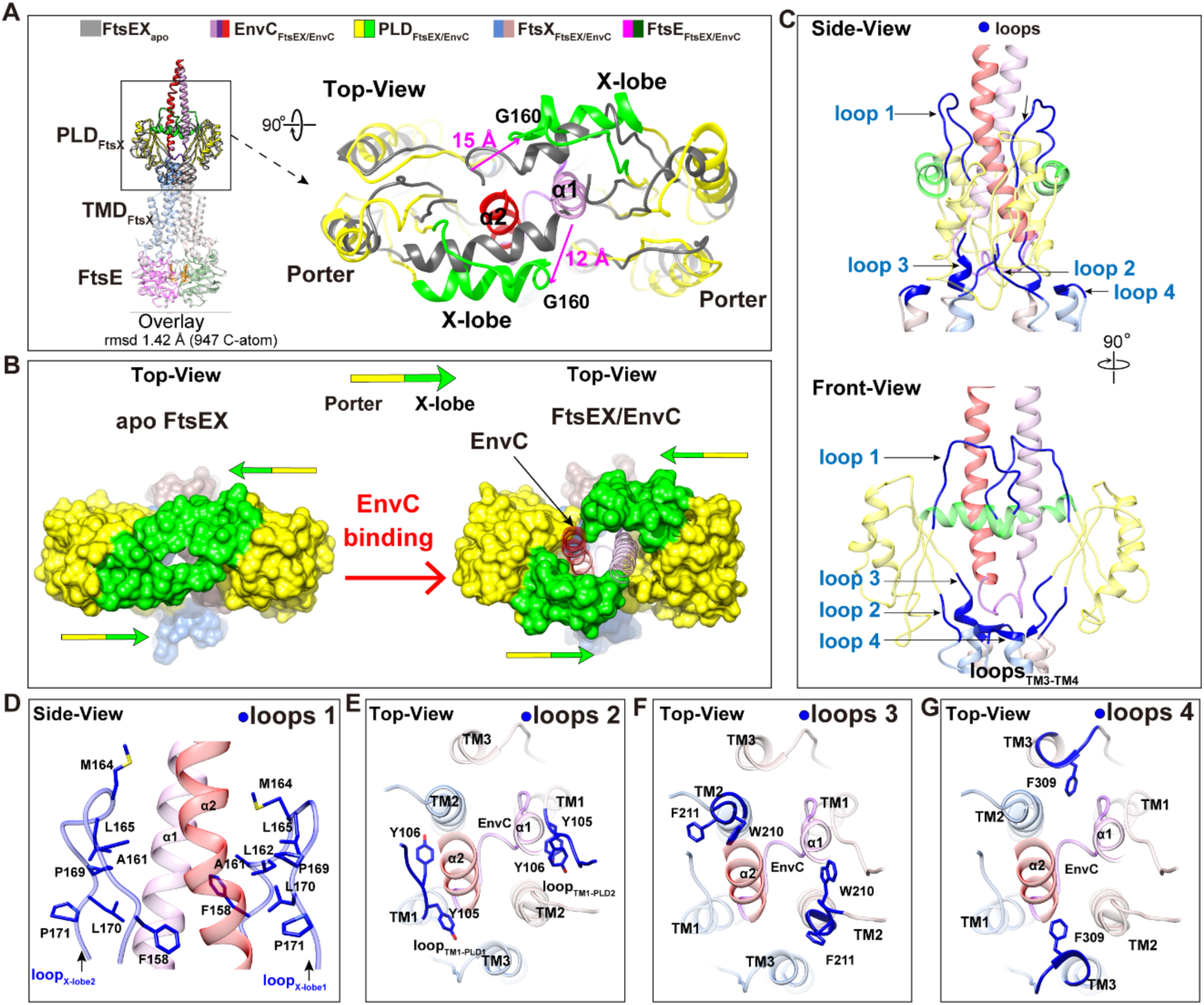
Identification of key loops for EnvC binding in the *Eco*FtsEX/EnvC Complex. (**A**) Structural overlay of full-length *Eco*FtsEX and *Eco*FtsEX/EnvC complexes obtained in this study, with a focus on the conformational changes in the PLD domain induced by EnvC binding. The overall overlay of *Eco*FtsEX and *Eco*FtsEX/EnvC is depicted in ribbon form. Color scheme: FtsE (magenta and dark green), FtsX (cornflower blue and rosy brown), Porter of PLD_FtsX_ (yellow), and X-lobe of PLD_FtsX_ (green). The TMD and FtsE regions are set at 60% transparency. (**B**) Depiction of the conformational changes in the PLD domains upon EnvC binding. (**C**) Side and front views illustrating the key loops involved in EnvC binding within the *Eco*FtsEX/EnvC complex. Key loops of FtsX, highlighted in blue, are indicated by arrows and labeled. (**D**) Side-view displaying the binding details between EnvC and flexible loops located in the X-lobe (G157-L171). (**E**), (**F**)**, and** (**G**) Top-down views demonstrating the binding details of EnvC and flexible loops positioned between TM1 and PLD (Y105-Q109) (**E**), PLD and TM2 (D208-L214) (**F**), and TM3 and TM4 (Q307-T311) (**G**).

Upon detailed examination, we noticed that the motifs in FtsEX directly interacting with EnvC are primarily loops (**Figure 4C**). The main loop engaging with EnvC is X-lobe (**Figure 4D**), covering residues G157-L171. This segment, originally disordered in the FtsEX alone structure, becomes crucial for engaging with EnvC, involving a binding interface of about 1204 Å^2^. The other interacting regions are loops spanning residues Y104-Q109 (**Figure 4E**) and D208-L214 (**Figure 4F**), and a segment of the small loop in the PLD domain (Q307-T311) (**Figure 4G**). Combined, these loops (eight in total from the two PLD domains) are responsible for the majority of the interactions with EnvC. This inherent adaptability in the loops provides FtsEX the versatility needed for binding the asymmetrical EnvC. Notably, the interactions contributed by these loops are largely hydrophobic (**Figure 4D-G**). Thus, our data suggest that the flexible loops, along with the hydrophobic interactions they facilitate, equip the symmetrical FtsEX to effectively engage with asymmetrical partners.

### Reconstitution and Elucidating the Elongated Conformation of *E. coli* FtsEX/EnvC/AmiB Complex for PG cleavage

The absence of the LytM density, in comparison to the crystal structure of PLD/EnvC (22), indicates that our ATP-bound FtsEX/EnvC sample likely has the LytM domain released, primed for hydrolase interaction. To further validate this, we conducted pull-down assays (**Figure 5A**). When we incubated AmiB with either the FtsEX/EnvC complex or the mutant FtsEX^E163Q^/EnvC in the presence of ATP or its nonhydrolyzable analogue, ATPγS, both the wild type and mutant FtsEX/EnvC complexes effectively pulled down AmiB. Notably, ATP and ATPγS showed similar levels of AmiB binding, confirming that ATP-bound FtsEX/EnvC is prepped for AmiB recruitment. Additionally, we investigated the influence of AmiB binding on FtsEX’s ATPase activity (**Figure 5B**). There was no obvious effect upon the addition of AmiB to the FtsEX/EnvC complex, suggesting that AmiB does not regulate FtsEX function.

**Figure 5.**
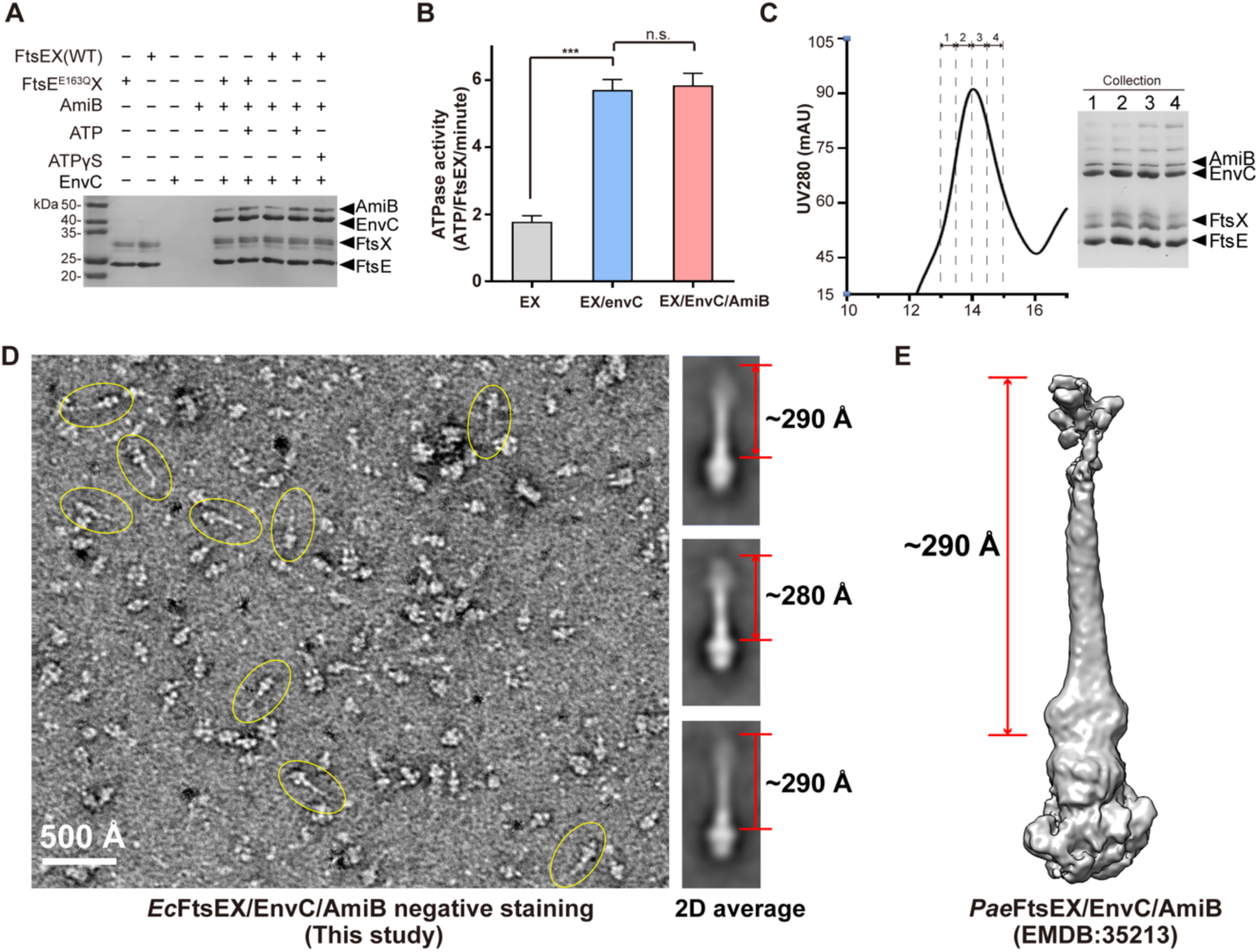
Biochemical reconstitution and negative-staining EM study of the AmiB-Bound *Eco*FtsEX/EnvC Complex. (**A**) Pull-down study illustrating the interaction between AmiB and FtsEX-EnvC or its ATPase mutant, in the presence or absence of ATP or its analogs. The initial FtsEX or FtsE^E163Q^X added to the resin contains 2 mM ATP, the ATP was subsequently removed by the wash buffer. (**B**) Evaluation of ATPase activity in FtsEX, FtsEX/EnvC, and FtsEX/EnvC/AmiB complexes. Notably, the binding of AmiB does not significantly impact the ATPase activity of FtsEX/EnvC. Data represent an average of three replicates, with error bars indicating mean ± SD. Statistical analysis performed using a two-tailed unpaired t-test; ***P < 0.0005. (**C**) SEC profile and SDS-PAGE gel of purified FtsEX/EnvC/AmiB complex reconstituted in peptidiscs. (**D**) Negative-staining micrograph and 2D average image of *Eco*FtsEX/EnvC/AmiB. (**E**) Cryo-EM density map of *Pae*FtsEX/EnvC/AmiB.

Next, we sought to check if the bound AmiB is activated for PG cleavage. FtsEX/EnvC complex was first mixed with a threefold molar excess of AmiB in the presence of 2 mM ATP, and the mixture was applied to size-exclusion chromatography (SEC) (**Figure 5C**). An early eluting fraction was observed, and subsequent SDS-PAGE confirmed the co-existence of FtsE, FtsX, EnvC, and AmiB in this fraction. The reconstituted PG-cleaving complex of *Eco*FtsEX/EnvC/AmiB, was further analysed using negative staining (NS) electron microscopy (**Figure 5D**). Many of the observed particles exhibited long tails in the image, a feature suggestive of AmiB binding. A 2D classification analysis presented an elongated structure, mirroring previous observations in *P. aeruginosa* (26). The electron densities on the periplasmic side displayed a distinct rod-like form in alignment with the FtsEX complex’s central axis, extending about 290 Å into the periplasm, which we attribute to the bound EnvC and AmiB. This conformation likely represents an activated state prepared for PG cleavage. However, due to the *Eco*FtsEX complex’s instability during the PG hydrolysis assay, we couldn’t directly check PG cleavage activity. The attempt to obtain a CryoEM structure of the complex was also unsuccessful, possibly because of its inherent flexibility, preventing the reconstruction of a clear EM map. Nonetheless, it has been well established in *P. aeruginosa*, where the active FtsEX/EnvC/AmiB complex also assumes an extended form (**Figure 5E**), we propose that our observed complex with a similar conformation is also in an activated state, primed for PG degradation.

## Discussion

A structural and functional description is here provided for the complete *E. coli* lytic machinery regulated by the FtsEX system. We observed an antiparallel arrangement of PLDs in the periplasmic space in the *Eco*FtsEX complex. Incorporation of ATP, but not hydrolysis, is capable of facilitating EnvC binding. The ternary complex increases its ATPase activity, compared with that of the binary complex. Interestingly, upon ATP binding, the flexibility of loops in the PLD domain allows a symmetric interaction with the EnvC coiled-coil domain that is compressed to allow the release of its LytM domain. While resolution for the complete FtsEX/EnvC/AmiB complex limited detailed descriptions, our cryo-EM data points to a very similar disposition of the lytic complex to that reported for *P. aeruginosa* (26) with a fully expanded EnvC coiled-coil that would place the amidase at ∼300 Å over the inner membrane.

In the context of recruiting an asymmetric partner by the inherently symmetrical FtsEX complex, our study has unveiled a dynamic and intriguing mechanism. At the centre of this mechanism lies the highly flexible PLD domain (composed of the porter domain and the loops connecting the different TM of FtsX), which demonstrates exceptional adaptability due to the presence of multiple flexible loops, including the x-lobe. These flexible loops within the PLD domain function as dynamic connectors, enabling the FtsEX complex to finely tune its conformation and engage with asymmetric partners in an exceptionally adaptable manner. This inherent plasticity empowers the FtsEX complex to align itself with the specific structural requirements of its binding partners, thereby facilitating their effective interaction and subsequent regulatory processes.

Moreover, these flexible elements serve a dual role by providing the necessary adaptability to bridge the gap between the inherent symmetry of the FtsEX complex and the asymmetry present in its interacting partners. This becomes particularly intriguing in the context of bacterial cell division, where the same FtsEX complex must interact with a variety of asymmetric binding partners. These binding partners exhibit diverse structural features and distinct binding interfaces, necessitating a mechanism that allows the inherently symmetrical FtsEX complex to engage with them with precision and effectiveness.

Our study has uncovered a previously unrecognized role of ATP in the stabilization of the FtsEX complex, a role that extends beyond its established function as a substrate. This discovery not only expands our understanding of the functional versatility of ATP but also unravels a crucial aspect of how FtsEX complexes are regulated. The stabilizing influence of ATP is particularly intriguing, as evidenced by our experiments across diverse bacterial species, suggesting that ATP-driven stabilization might represent a shared feature among FtsEX complexes in various bacteria. Furthermore, it is noteworthy that both the FtsEX systems from *E. coli* and the recently characterized system from *P. aeruginosa* exhibit a relatively low basal ATPase activity. However, this activity is significantly enhanced in the presence of their respective cognate binding partner, EnvC. Specifically, for *E. coli*, the ATPase activity increases by a factor of 2.2, whereas for *P. aeruginosa*, the increase is even more pronounced, with a greater than 20-fold enhancement observed. This additional regulatory mechanism ensures that, under normal physiological conditions, minimal energy is expended. It is only during the cell division process, when components like EnvC are involved, that the ATPase activity of FtsEX becomes fully activated.

Interestingly, our cryoEM map of *Eco*FtsEX, expressed from its native host bacteria *E. coli*, revealed an additional density located on the bottom side of the complex adjacent to FtsE (**Figure supplement 2**, and **Figure supplement 10A**). This suggests the co-purification of endogenously binding proteins with the complex. To identify the proteins, we conducted mass spectrometry on the purified FtsEX sample. Our results showed that in addition to FtsE and FtsX, FtsA was present in significant abundance, followed by FtsZ (**Figure supplement 10B**). This observation is consistent with previous genetic and phenotypic observations on the *E. coli* divisome activation (17, 27). We also noted during the course of these studies that the interaction of *E.coli* FtsA with FtsEX protein complexes from other species is indicative of the conserved and fundamental nature of divisome protein interactions. For example, the overexpression of *S. pneumoniae* FtsEX in *E. coli* also indicated an association with *E.coli* FtsA and could be isolated using styrene-maleic acid isolation and affinity purification as shown in **Figure supplement 10C** (31, 32). If this holds true, it would provide a mechanism for the temporal regulation of PG cleavage at the cell division site, as FtsA and FtsZ are the key components of the constriction ring at the septum site (15, 18, 27, 33–37). However, the identity of this density and the association of FtsEX with other Fts components for divisome formation warrant further investigation to fully elucidate the role of FtsEX in bacterial cell division.

In combination with previous studies on the inactivated EnvC (22) and AmiB activation by *Pae*FtsEX (26), we propose a working model that may be conserved across Gram-negative FtsEX systems (**Figure 6**). Firstly, physiological ATP stabilizes the formation of a functional FtsEX regulator with low basal ATP hydrolysis activity; In the periplasm, the inactive EnvC interact with the flexible PLD domain of FtsEX, which adopts a highly asymmetric conformation, as shown in the inactivated EnvC structure (22). Our study, along with previous findings, highlights the crucial role of ATP binding in stabilizing FtsEX, likely by promoting FtsE dimerization in the cytosol. Meanwhile, EnvC binding plays a key role in stimulating FtsEX ATPase activity. Although EnvC binding appears nucleotide-independent, the precise sequence of ATP binding and EnvC recruitment during cell division remains to be further explored. Next, upon full recognition by FtsEX, considering the presence of ATP in cell and a symmetrical conformation defined within the context of a full-length FtsEX complex, recruited EnvC is compelled to adopt a relatively symmetrical conformation, involving squeezing of the coiled-coil domain to facilitate the release of the LytM domain. Finally, the released LytM recruits amidases through direct interaction, consequently removing the blocking helix from the catalytic site and activating PG cleavage. Remarkably, expansion of the coiled-coil domain in EnvC would place the amidase domain ∼300 Å over the inner membrane of the Gram-negative bacteria (as here seen in *E. coli*, and also in *P. aeruginosa* (26)), compatible with the location of the septal PG (26) and avoiding damages into the non-septal cell wall. Throughout this process, ATP binding appears to be sufficient, with ATP hydrolysis potentially serving as a driving force for the release of EnvC and amidases from FtsEX, allowing them to be recycled to a normal ABC transporter. This transporter by itself may play additional roles, such as antibiotic resistance, as suggested by previous studies (22). For the temporal regulation of PG cleavage, the potential interaction between FtsEX and FtsA/FtsZ, as indicated in this study and previous genetic research (17, 27), could bridge the PG-cleaving complex with septum constriction ring, facilitating precise cell division at the septum site. This model provides a potential conserved mechanism for PG cleavage activation by FtsEX and illuminates its temporal regulation. Although further validation is required, our results offer valuable biochemical and structural insights that complement and elucidate decades of genetic research in *E. coli* and other Gram-negative bacteria.

**Figure 6.**
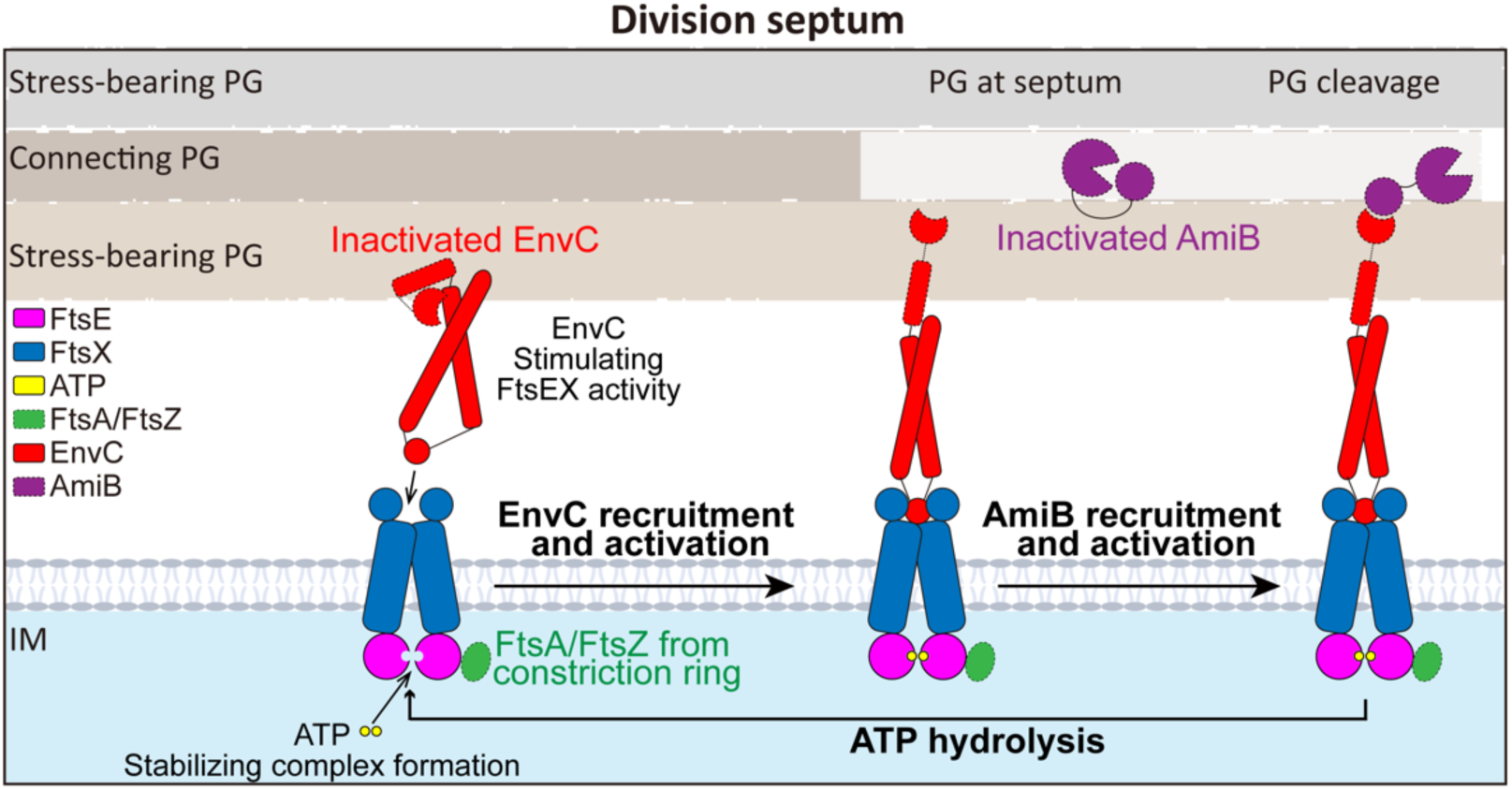
A working model illustrating FtsEX complex formation, recruitment of EnvC and AmiB, and the temporal regulation of PG cleavage activation in Gram-negative bacterial systems. Initially, in the cytosol, physiological ATP stabilizes the formation of the FtsEX complex, while in the periplasm, the asymmetrical and inactive EnvC interacts with the flexible PLD domain of FtsEX, stimulating its ATPase activity. However, the precise sequence of ATP binding and EnvC recruitment in cells remains under debate. Upon complete recognition by FtsEX, the CCD domain of recruited EnvC undergoes a conformational change to a symmetrical state, involving compression of the coiled-coil domain, facilitating the release of the LytM domain. Interaction with EnvC activates FtsEX’s ATPase activity. The liberated LytM recruits amidases through direct interaction with the catalytic domain’s blocking helices, leading to the removal of the obstructing helix from the catalytic site and activation of PG cleavage. ATP hydrolysis likely triggers the release of EnvC and AmiB, allowing FtsEX to return to its normal ABC transporter state with a low basal ATPase activity. The potential interaction between FtsEX and the constriction ring component FtsA/FtsZ precisely locates the PG-cleaving complex at the septum for temporal regulation of PG cleavage.

## Materials and Methods

### Cloning and Mutagenesis

The primers and templates utilized in this study are listed in **Table supplement 1**. To generate the PCR amplicon containing FtsEXs in genomic DNA of *Escherichia coli* Str *K-12 MG1655*, *Streptococcus pneumoniae str. D39*, *Bacillus_subtilis_subsp_subtilis_ATCC_23857*, and *Pseudomonas aeruginosa ATCC 47085D*. ClonExpress II One Step Cloning Kit (Vazyme) was used, followed by purification. After purification through gel electrophoresis, the resulting product was ligated to pTD68 vector (PMID: 20300061) digested with the Nde I and Hind III enzymes to produce pTD68 [PT7::his6-FtsEX]. The construction of pET28-sumo (PT7:his6-sumo-*Eco*EnvCΔ34), (PT7:his6-sumo-*Eco*AmiBΔ22), involved ligating pET28-sumo digested with BamH I and Xho I to a PCR amplicon synthesized by EnvC-F, EnvC-R, AmiB-F, AmiB-R which were also digested with the same enzymes. The plasmids utilized in this study are listed in **Table supplement 2.** Mutations to *Eco*FtsE was generated utilizing the QuikChange kit (Agilent Technologies), using pTD68 and pET28-sumo as templates and the primers listed in **Table supplement 1**. The mutations were then verified by DNA sequencing.

### Protein expression, purification of FtsEX complex and their mutants

FtsEX and its mutants are expressed in *E. coli* strain BL21(DE3) pLysS (Invitrogen) using lysogeny broth (LB) supplemented with 50 µg/ml ampicillin and 10 µg/ml chloramphenicol. This is incubated at 37°C until an optical density at 600 nm (OD_600_) of 0.8 is achieved. Following this, induction with 0.5 mM isopropyl-β-D-thiogalactoside (IPTG) takes place at 20°C for 16 hours. The cells are subsequently harvested through centrifugation using a Beckman JLA9-1000 rotor at 5,180 xg and 20°C for 15 minutes. Harvested cells are then stored at −80°C.

For membrane extraction, the frozen cell pellets undergo thawing at 4°C and are resuspended in buffer A, consisting of 50 mM Tris-HCl pH 8.0, 250 mM NaCl, 10% glycerol, and 2 mM ATP/MgCl_2_. Lysis of the mixture was achieved with a high-pressure homogenizer (AH-100D, ATS). Subsequent centrifugation was performed at 27,000 xg for an hour at 4°C. Following this, solubilization of the membrane takes place in buffer B, containing 50 mM Tris-HCl, 250 mM NaCl, 1.3% n-dodecyl β-D-maltoside (DDM), pH 8.0, 10% glycerol, and 2 mM ATP/MgCl_2_, and agitated for 2 hours at 4°C. Post a 30-minute centrifugation at 100,000 xg, the solubilized membrane fraction gets incubated with TALON SuperFlow resin (Takara 635502) for 1 hours at 4°C and was later washed using buffer B supplemented with 20 mM imidazole. The present detergent is substituted by a peptide (GenScript) to form peptidiscs, employing the “on-beads” method (38, 39) with a final peptide concentration of 1mg/ml. The FtsEX complex, reconstituted in the peptidiscs, is washed using buffer B inclusive of 20 mM imidazole and eluted with buffer B rich in 200 mM imidazole. The purified protein is directed through a gel filtration chromatography column (Superose 6 increase 10/300) and is eluted with buffer C, which includes 25 mM Tris-HCl, 150 mM NaCl, pH 8.0, and 2 mM ATP/MgCl_2_. The fractions bearing the FtsEX complex are combined together, concentrated to 3 mg/ml by concentrator (Amicon® Ulyra-15 100kDa), and preserved at −80°C for further protein interaction, pull-down, and ATP hydrolysis assay.

### Protein expression, purification of EnvC, AmiB

For the expression of EnvC, AmiB and their respective variants, the *E. coli* strain BL21(DE3) pLysS was employed. These cells were cultured in LB medium with 50 µg/ml kanamycin at 37°C. Once the OD_600_ reached a value of 0.6, induction was facilitated by the addition of IPTG to achieve a final concentration of 0.5 mM, and this was maintained for 16 hours at 20°C. Following the induction, cells were harvested through centrifugation using a JLA9-1000 rotor in a Beckman centrifuge at 5,000 rpm and 20°C for a duration of 15 minutes. The resulting pellets were stored at −80°C. For further processing, these frozen pellets were brought to 4°C for thawing, using buffer C (50 mM Tris-HCl pH 8.0, 150 mM NaCl, and 10% glycerol) and then subjected to lysis via a high-pressure homogenizer (AH-100D, ATS). Subsequent centrifugation was performed at 27,000 xg for an hour at 4°C. The resultant supernatant was incubated with TALON resin, followed by a wash using buffer C complemented with 20 mM imidazole. Post-washing, the His-tagged SUMO protease Ulp1 was introduced to the resin-buffer at a ratio of 0.1 mg Ulp1 for every liter of culture, facilitating the cleavage of the SUMO tag over 2 hours at ambient temperature. The processed samples were isolated, concentrated, and then passed through the Superose 6 increase 10/300 column previously balanced in buffer D which includes 25 mM Tris-HCl, 150 mM NaCl, pH 8.0. Target protein was confirmed using SDS-PAGE, further concentrated up to 3 mg/ml, and stored at −80°C.

### Analyzing the Impact of ATP on the FtsEX Complex

Expression and purification of FtsEXs which from *Escherichia coli (Eco)*, *Streptococcus pneumoniae (Sp)*, *Bacillus_subtilis (Bs)*, and *Pseudomonas aeruginosa (Pae)* followed the procedure above. For the ATP-free FstEXs purification, the whole process excludes ATP. For the samples of FtsEXs with ATP or ATP analogs, 2 mM ATP or ATP analogs present in the purification process. The processed samples of *Sp*FtsEX, *Bs*FtsEX, *Pae*FtsEX were concentrated, and then passed through the Superose 6 increase 10/300 column previously balanced in buffer which includes 25 mM Tris-HCl, 150 mM NaCl, pH 8.0 with or without 2 mM ATP. The eluted proteins profiles were confirmed using SDS-PAGE gel. For the *Eco*FtsEX/EnvC samples purification add EnvC with final concentration 1mg/ml after FtsEX reconstituted in the peptidiscs. The eluted ATP-free *Eco*FtsEX sample was performed at 12,000 xg for 15 min at 4°C due to persistent precipitate appearing in the eluted sample. Confirming the supernatant and precipitate using SDS-PAGE gel.

### Protein interaction assay

To explore the interaction between FtsEX and EnvC, we employed affinity chromatography with TALON resin. Specifically, 15 μg of FtsEX or its ATPase-defective variants were introduced to the resin pre-equilibrated at room temperature, utilizing a Micro Gravity Column, for a 15-minute duration. After washing away unbound FtsEX with 100 μl of buffer E (50 mM Tris-HCl pH 8.0, 150 mM NaCl, 10% glycerol, and 2 mM MgCl_2_) supplemented with 5 mM imidazole (to get rid of the extra ATP present in the protein), we incorporated 5 μg of EnvC diluted in 60 μl of the aforementioned buffer in the presence and absence of ATP or ATP analogs. This mixture was incubated for an additional 10 minutes. Subsequently, the beads were rinsed with buffer E supplemented with 5 mM imidazole in the presence and absence of ATP or ATP analogs. To elute the bound proteins and possible interaction partners, we utilized 20 μl of buffer E with a concentration of 300 mM imidazole, either with or without 2 mM ATP or ATP analogs. Eluted proteins were then analyzed using SDS-PAGE.

To explore the interactions between FtsEX/EnvC and AmiB, 20 μg of FtsEX/EnvC or its ATPase-deficient variants, FtsE^E163Q^X/EnvC, were added to the TALON resin. This was done either in the presence or absence of 2 mM ATP or ATP analogs and incubated at room temperature for 15 minutes. Any free FtsEX/EnvC or FtsE^E163Q^X/EnvC was removed by washing the resin with 100 μl of buffer E containing 5 mM imidazole. Following this, 10 μg of AmiB were introduced to the resin with or without 2 mM ATP or ATP analogs. After a 10-minute incubation, a triple wash was executed using 100 μl of buffer E supplemented with 5 mM imidazole, either with or without 2 mM ATP or ATP analogs. Finally, bound proteins were eluted using 20 μl of buffer E supplemented with 300 mM imidazole, either with or without 2 mM ATP or ATP analogs, and the elution profile was assessed via SDS-PAGE.

### Reconstitution of FtsEX complexes

For the reconstitution of the FtsEX-EnvC complex, an excess of EnvC, three times the amount of FtsEX, was introduced to the resin-bound FtsEX-peptidiscs during the purification process. This mixture was allowed to interact at room temperature for 15 minutes. Subsequent to this incubation, any free EnvC was removed by washing with buffer A containing 20 mM imidazole. The FtsEX-EnvC complex was subsequently eluted from the resin using buffer A supplemented with 200 mM imidazole. This eluted protein was applied to additional purification through gel filtration on a Superose 6 increase 10/300 column, equilibrated in buffer B. Fractions containing the FtsEX-EnvC complex were assessed using SDS-PAGE, combined, concentrated to a concentration of 3 mg/ml, and stored at −80°C.

For the reconstitution of the AmiB-EnvC-FtsEX supercomplex, AmiB was mixed with the FtsEX-EnvC complex in peptidiscs, at a 3:1 ratio. Following a 15-minute incubation, the supercomplex was further purified via gel filtration. Fractions encompassing the FtsEX-EnvC-AmiB complex were collected, concentrated to 3 mg/ml, and subsequently stored at −80°C.

### ATP hydrolysis assay

To assess the ATPase activity of FtsEX and its mutants or complxes, we employed the malachite green phosphate assay (40). We introduced 2 μg of FtsEX with or without 0.7 μg of EnvC or 0.7 μg AmiB of following 0.8 μg of EnvC into a 20 μl reaction buffer comprising 25 mM Tris-HCl (pH 8.0), 150 mM NaCl, and 2 mM ATP/MgCl_2_. For the elucidation of the Michaelis-Menten constants, varying concentrations of ATP/MgCl_2_, ranging from 0.05 to 5 mM, were utilized to initiate the enzymatic reaction. After a 10-minute incubation at 37°C, 80 μl of the Malachite Green-Ammonium Molybdate (MG-AG) solution was added and the resultant mixture was quickly vortexed to deactivate the enzyme. A brief 2-minute incubation was followed by the addition of 10 μl of a 34% (w/v) sodium citrate solution, with a subsequent vortexing step. Absorbance was then recorded at 650 nm after a 5-minute interval. Quantification of released inorganic phosphate (Pi) was achieved using a calibration curve derived from known Pi concentrations in a K_2_HPO_4_ solution. Data processing and curve fitting to the Michaelis-Menten equation were conducted utilizing Microsoft Excel in conjunction with GraphPad Prism 9.

### Negative staining imaging of FtsE^E163Q^X-EnvC-AmiB complex

For negative staining of the FtsE^E163Q^X-EnvC-AmiB sample, it was diluted to a concentration of 0.01 mg/ml. A 5 μl aliquot of the sample was deposited onto a 300 mesh copper grid coated with carbon film (EMS). After allowing the sample to interact with the grid for 60 seconds, any surplus was carefully blotted away using filter paper. The grid was then washed with 5 μl of Uranyless EM Stain (EMS). Subsequent to the washing step, the grid was stained by applying another 5 μl of the Uranyless EM Stain for 120 seconds. Excess stain was again removed with filter paper. The prepared grid was left to air-dry under ambient conditions for approximately 15 minutes. Imaging of the sample was executed with a Tecnai 12 microscope (FEI), operating at 120kV with a LaB6 source, and magnification set to 52, 000 x.

### Cryo-Electron microscopy sample preparation and data acquisition

To prepare the sample, 3.5 μl of the protein sample at a concentration of 3 mg/ml was applied to Quantifoil holey carbon grids (R1.2/1.3, 400 mesh) that were glow-discharged. To obtain the ATPγS -bound complex, the sample was incubated at 37°C for 8 minutes with 3mM ATPγS-2mM Mg^2+^ (FtsEX) or 2 mM ATP-Mg^2+^ (FtsE^E163Q^X/EnvC) before freezing. The grids were blotted for 3.5-4 seconds with 100% relative humidity and then plunge-frozen in liquid ethane that was cooled by liquid nitrogen using a Vitrobot System (Gatan).

Cryo-electron microscopy data was collected at liquid nitrogen temperature using a Titan Krios electron microscope (Thermo Fisher Scientific), equipped with a K3 Summit direct electron detector (Gatan) and GIF Quantum energy filter. All cryo-EM movies were recorded in counting mode with SerialEM4 (41) with a slit width of 20 eV from the energy filter. Movies were acquired at nominal magnifications of 81k x, which corresponded to a calibrated pixel size of 1.06 Å on the specimen level. The total exposure time for each movie was 5 seconds, resulting in a total dose range of 35-39 electrons per Å^2^, fractionated into 34 frames. Additional electron microscopy data collection parameters can be found in **Table supplement 3**.

### Cryo-Electron microscope image processing

For electron microscopy (EM) data processing, CryoSPARC (42) was utilized. Dose-fractionated movies, captured with a K3 Summit direct electron detector, was first applied to motion correction using MotionCor2 (43). A sum of all frames from each movie was generated applying a dose-weighting scheme. This sum was utilized for all subsequent image-processing steps, except for defocus determination. For that, CTFFIND4 (44) was used on the non-dose-weighted summed images from all movie frames. The initial particle selection was achieved using the blob picker, which was subsequently refined by the template picker. The steps of 2D and 3D classification, as well as 3D refinement, were carried out using the “2D Classification”, “Ab-initial Reconstruction”, and “Heterogeneous Refinement” functions. For further refinement, both the “Homogeneous Refinement” and “Non-Uniform Refinement” functions were employed. The overall resolutions was determined using the gold-standard Fourier shell correlation (FSC) threshold of 0.143. Lastly, the local resolution was assessed via the “Local Resolution Estimation” tool.

### Model building and refinement for FtsEX and its complex with EnvC

The cryo-EM maps of the ATPγS-bound FtsEX and ATP-bound FtsE^E163Q^X/EnvC complexes were resolved at resolutions of 3.7 Å and 3.4 Å, respectively, with sufficient quality to enable accurate model building for the TMD domain, FtsE, and a portion of the PLD region (see **Figure supplement 3** and **Figure supplement 6** for details), leaving only the N-terminal region of FtsX, including residues 1-52, not modeled.

Initial models for FtsE and FtsX were generated using the AlphaFold program (42). Subsequently, the FtsEX models were rigid-body fitted into the cryo-EM map density using UCSF Chimera software (45). These initial models were then refined manually within Coot (46). Restraints for ATPγS were generated using the phenix_elbow program, employing its isomeric SMILES string files obtained from the PDB Chemical Component Dictionary via Ligand Expo. Further refinement was performed using Phenix (47), including ‘minimization_global’ and ‘Real-space refinement’ protocols. Multiple rounds of manual adjustments in Coot, in conjunction with real-space refinement in Phenix, were carried out to obtain the final model. Model validation was performed using Ramachandran plots, MolProbity scores, and Clash scores in Phenix (47) (details provided in **Table supplement 3**).

In the case of the FtsE^E163Q^X/EnvC/ATP complex structure, the initial models for EnvC was derived from AlphaFold (48) and our resolved FtsEX structure was used as initial model for FtsEX part. These initial models were fitted into the cryo-EM map of the FtsE^E163Q^X/EnvC/ATP complex using UCSF Chimera (45). Subsequent manual adjustments were performed within Coot (46), and the Glu163 was replaced with Gln. Similar to the ATPγS-containing complex, restraints for ATP were generated using the phenix_elbow program. The model was refined using phenix software (47), incorporating ‘minimization_global’ and ‘Real-space refinement’ procedures. The final model was refined through multiple iterations of manual adjustments in Coot (46), alongside real-space refinement in Phenix. Model quality assessment included Ramachandran plots, MolProbity scores, and Clash scores, with detailed results provided in **Table supplement 3** (**47**).

## Author Contributions

Jianwei Li and Yutong He performed biochemical reconstitution, functional characterization, and cryo-EM studies; Xin Xu performed kinetics assays; Souvik Naskar and Nicholas S. Briggs conducted co-purification studies of endogenous FtsA/FtsZ with FtsEX in *E. coli*. Martin Alcorlo, David Roper, Juan A. Hermoso and Lok-To Sham helped with data analysis; Jian Shi helped with EM data collection; Min Luo designed and supervised research, analyzed data, and wrote the paper with help from all authors.

## Acknowledgments

We appreciate the contributions of Yibei Xiao and Zhaoxing Li from China Pharmaceutical University, whose expertise in Mass Spectrometry greatly enhanced our research. Furthermore, we thank all Luo lab members for their constructive comments throughout the course of this study.

## Data and materials availability

Two three-dimensional cryo-EM density maps of FtsEX and its complexes with EnvC have been deposited in the Electron Microscopy Data Bank under accession codes: EMD-37324 (ATPγS-bound FtsEX); EMD-37325(ATP-bound FtsE^E163Q^X with EnvC). Two atomic models have been deposited in the Protein Data Bank under accession codes 8W6I (ATPγS bound FtsEX), and 8W6J (ATP-bound FtsE^E163Q^X/EnvC). All data presented in this study are provided within the main text and supplementary materials, or readily accessible upon request.

