## Supplemental information for "Structural insights into peptidoglycan hydrolysis by the FtsEX system in *Escherichia coli* during cell division"

**This PDF file includes:**

Figures supplement 1-10
Tables supplement 1-3
SI References

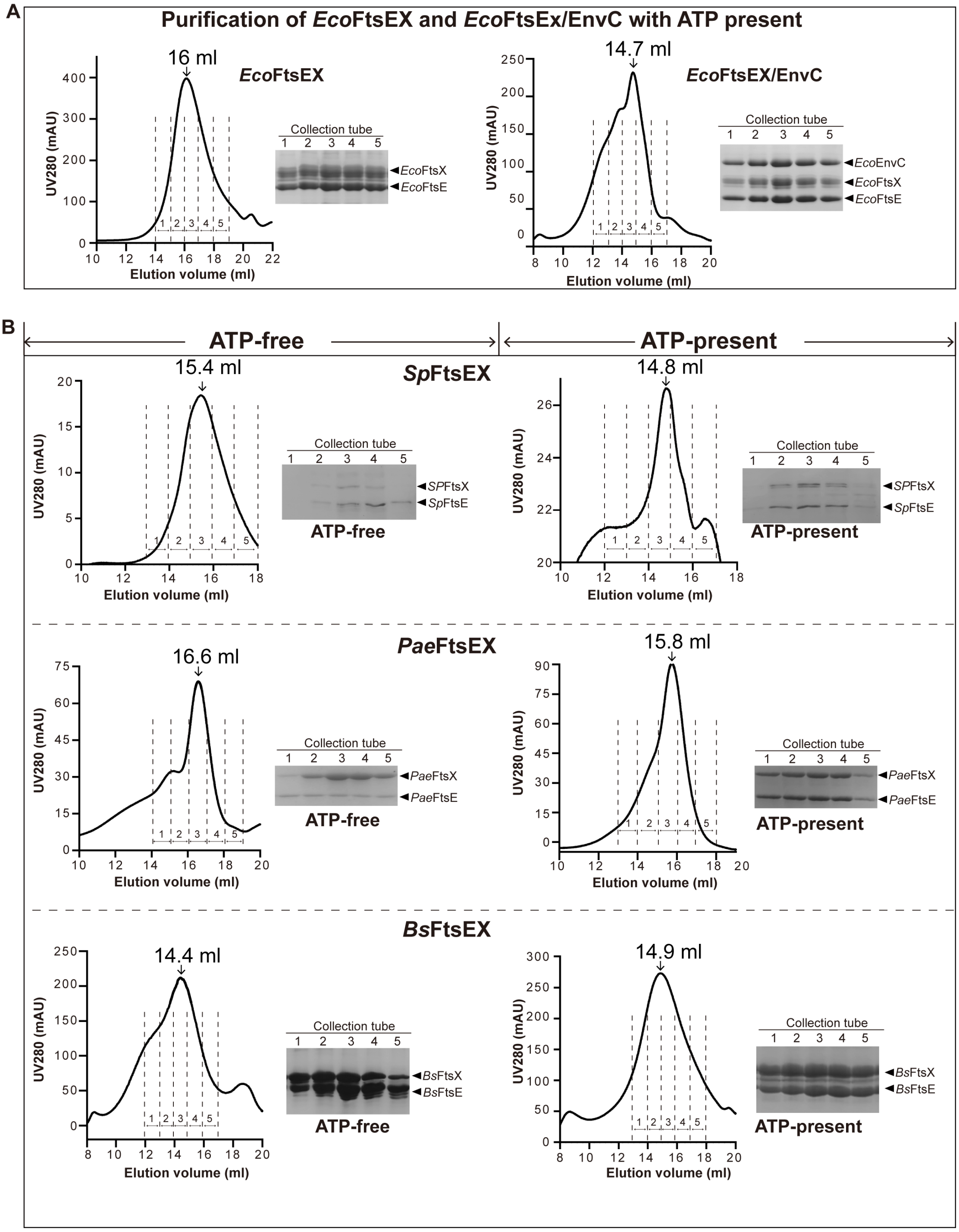


**Figure supplement 1. Size exclusion chromatography (SEC) analysis of the impact of ATP on the FtsEX complex. (A)** SEC profiles and SDS-PAGE gels depicting *Eco*FtsEX and *Eco*FtsEX/EnvC behavior in the presence of ATP. The inclusion of 2 mM ATP results in the formation of a stable complex for both *Eco*FtsEX and *Eco*FtsEX/EnvC, as demonstrated by their tight association throughout the SEC process. **(B)** SEC profiles and SDS-PAGE gels showcasing FtsEX complexes from *Streptococcus pneumoniae* (*Sp*), *Pseudomonas aeruginosa* (*Pae*), and *Bacillus subtilis* (*Bs*) in the presence or absence of ATP. Without ATP, FtsEX complexes tend to dissociate, while in the presence of ATP, they form stable complexes with correct stoichiometry.

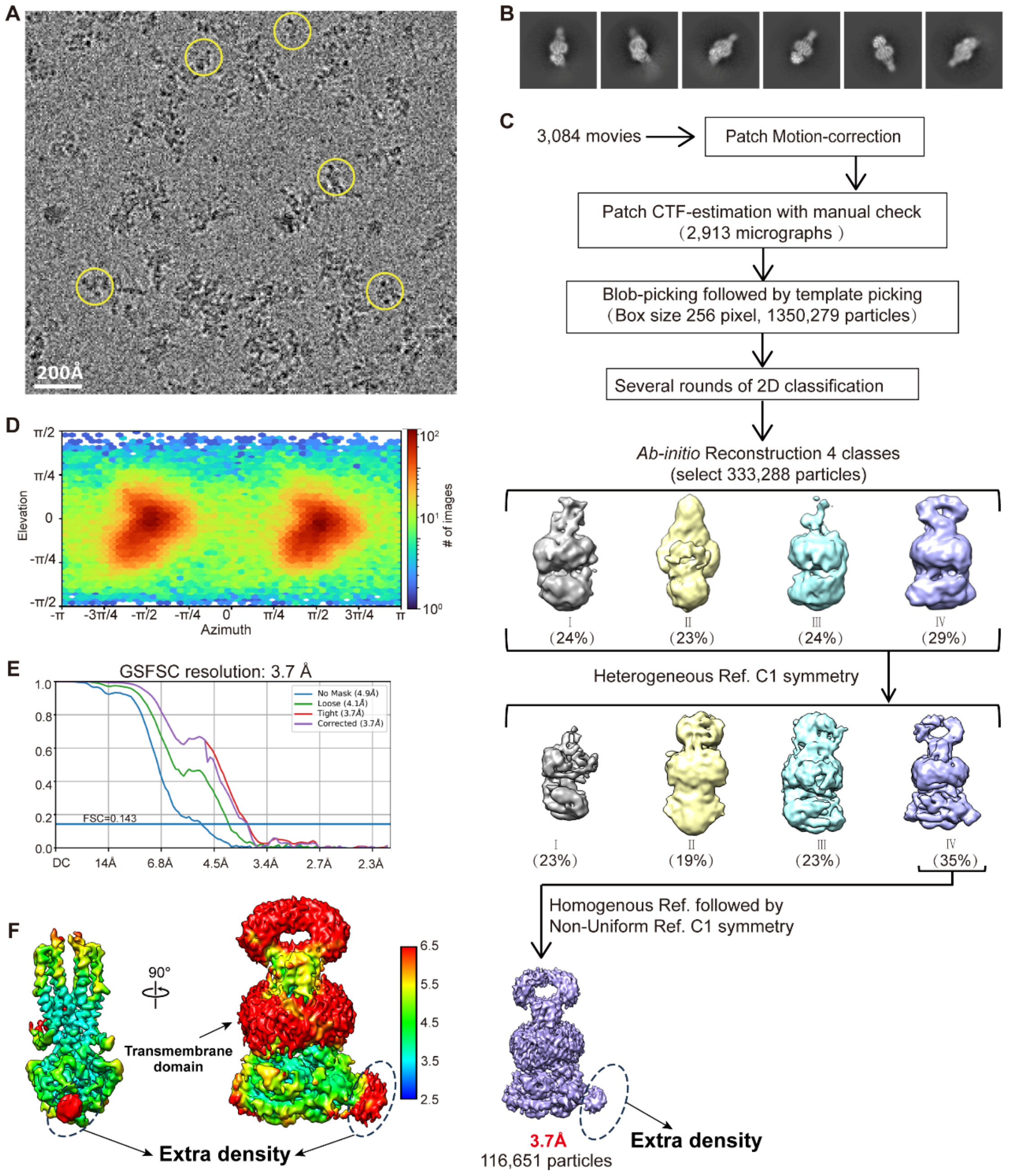


**Figure supplement 2. Biochemical reconstitution and single-particle cryo-EM analysis of FtsEX complex.** (**A**) Representative cryo-EM image with several particles boxed out by yellow circles. (**B**) Representative 2D averages of cryo-EM particle images. The box dimension is 270 Å. (**C**) Image processing flowchart. The final maps of one major conformation with its overall resolutions is indicated in red. The extra density uses dotted line to outline. (**D**) Angular distribution of the cryo-EM particles included in the final 3D reconstruction. (**E**) The Fourier shell correlation (FSC) curve: gold standard FSC between two half data maps with indicated resolution at FSC=0.143 (FSC corrected applied). (**F**) Side-view (contour level 0.22 in Chimera) and Front-view (contour level 0.1 in Chimera) show the surface cryo-EM map filtered to the estimated overall resolution and colored according to local resolution. The extra density use dotted line to outline.

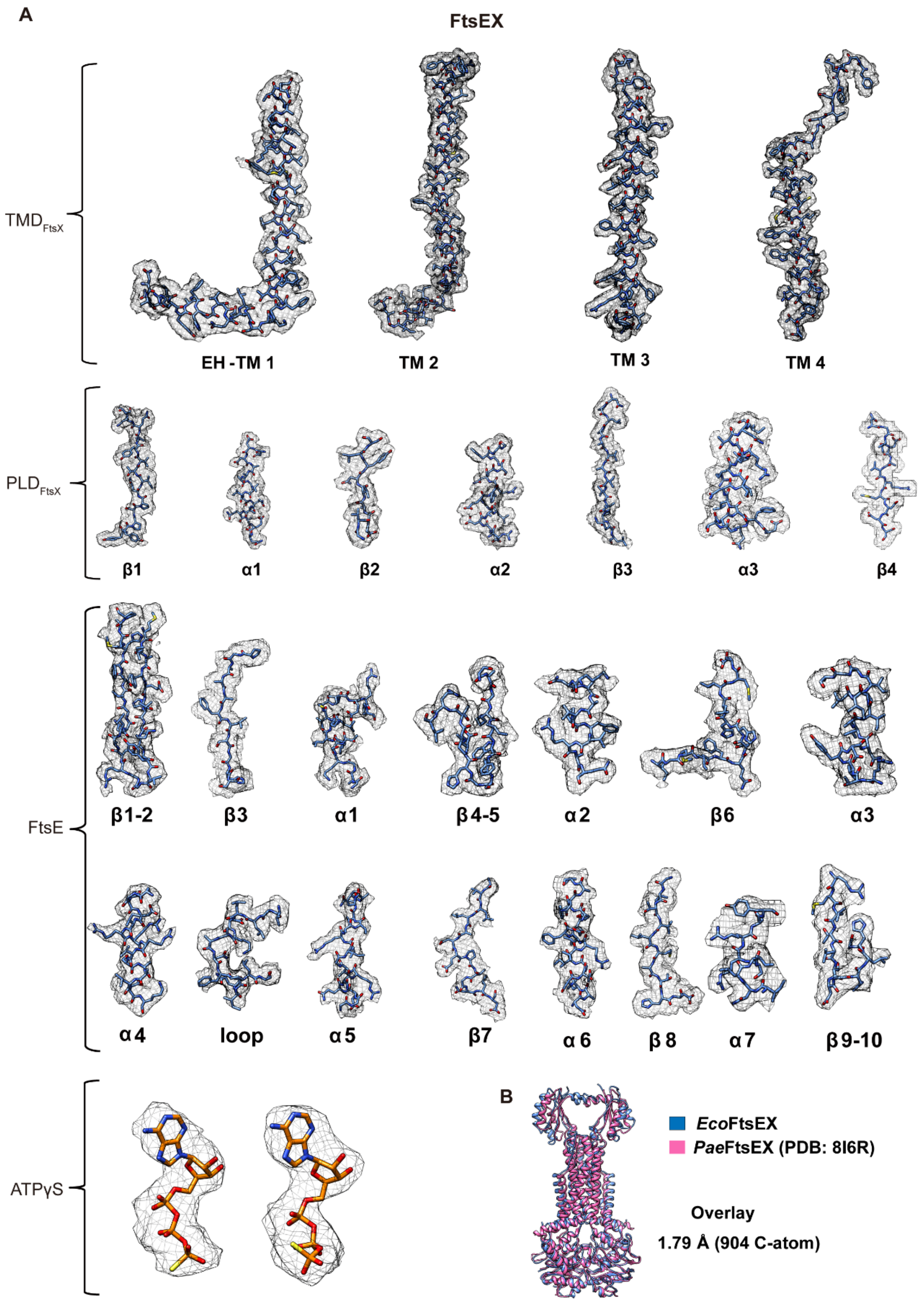


**Figure supplement 3. Cryo-EM density of different regions of FtsEX in the structure of FtsEX complex.** (**A**) Cryo-EM density of different regions of FtsE, FtsX and ATP in the structure of FtsEX complex. (**B**) Structural comparison of *Eco*FtsEX and *Pae*FtsEX (PDB: 8I6R), showing high structural similarity (1).

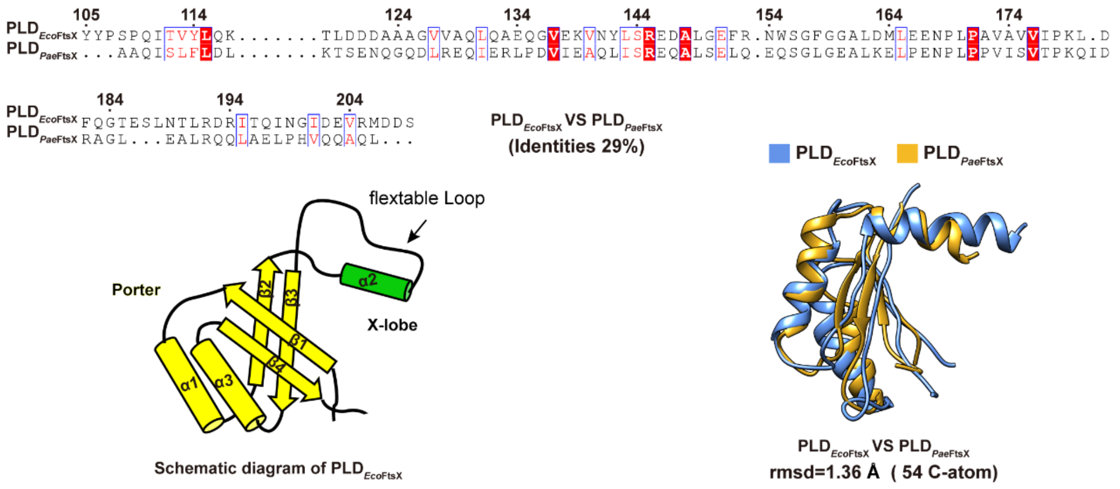


**Figure supplement 4. Structural comparison of PLD domain between *Eco*FtsEX and *Pae*FtsEX.** Sequence and structure alignment of PLD domain between *Eco*FtsEX and *Pae*FtsEX. Left: Topology diagram of PLD domain in *Eco*FtsEX. α helices are shown as cylinder, β sheets are shown as arrows, the Porter of $PLD_{FtsX}$ in yellow, and the X-lobe of $PLD_{FtsX}$ in green; Right: Structural comparison of PLD monomers between *Eco*FtsEX and *Pae*FtsEX with the structures overlaid on each other, the PLD of *Eco*FtsX in blue and the PLD of *Pae*FtsX in brown.

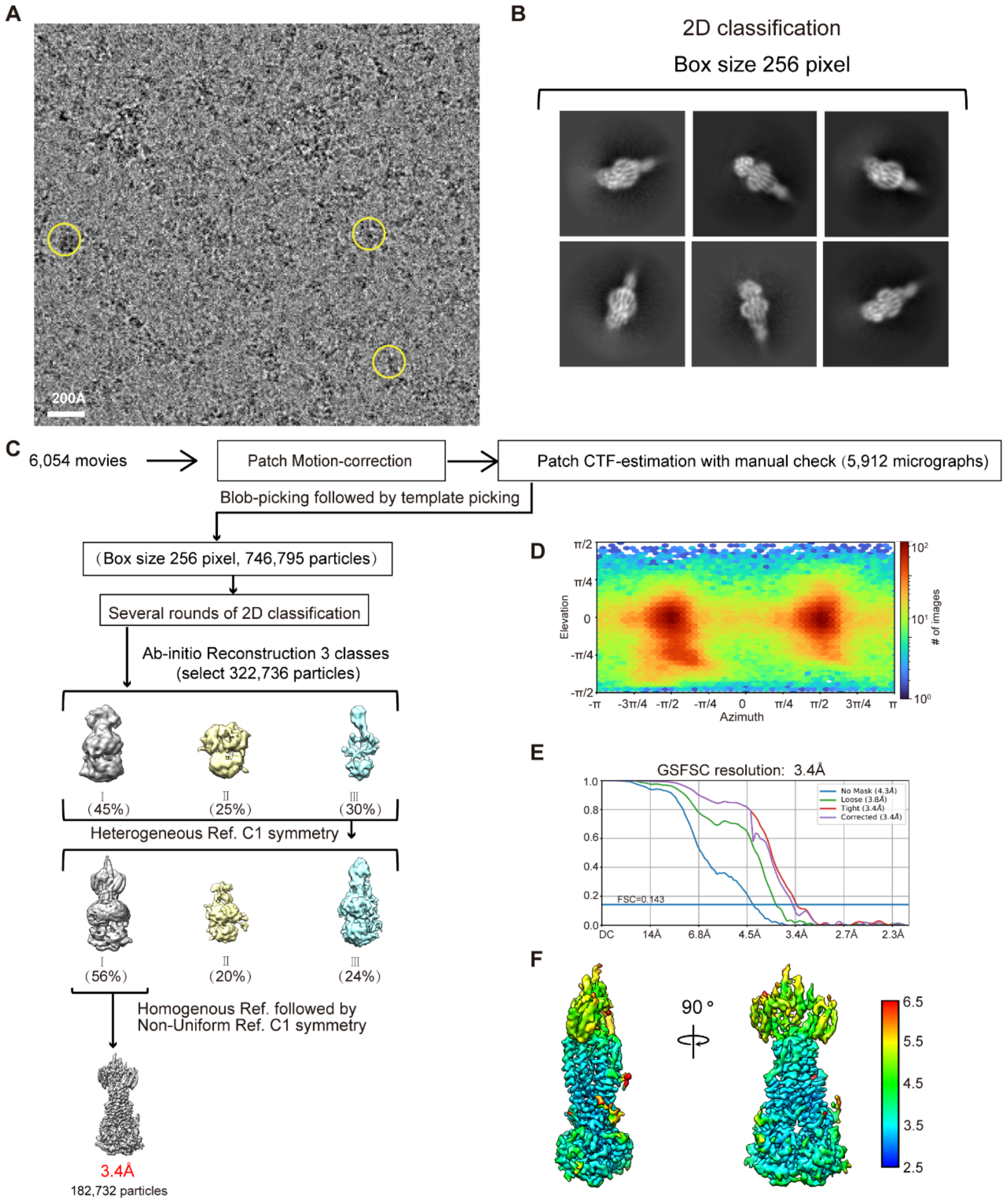


**Figure supplement 5. Biochemical reconstitution and single-particle cryo-EM analysis of FtsEX/EnvC complex.** (**A**) Representative cryo-EM image with particles marked by yellow circles (256 pixel box size). (**B**) Representative 2D averages of cryo-EM particle images with box dimension (270 Å). (**C**) Image processing flowchart with 256 pixel box size. The final maps of one major conformation with its overall resolutions is indicated in red. ((**D**)) Angular distribution of the cryo-EM particles included in the final 3D reconstruction. ((**E**)) The Fourier shell correlation (FSC) curve: gold standard FSC between two half data maps with indicated resolution at FSC=0.143 (FSC corrected applied). (**F**) The surface cryo-EM map filtered to the estimated overall resolution and colored according to local resolution (contour level: 0.19 in Chimera).

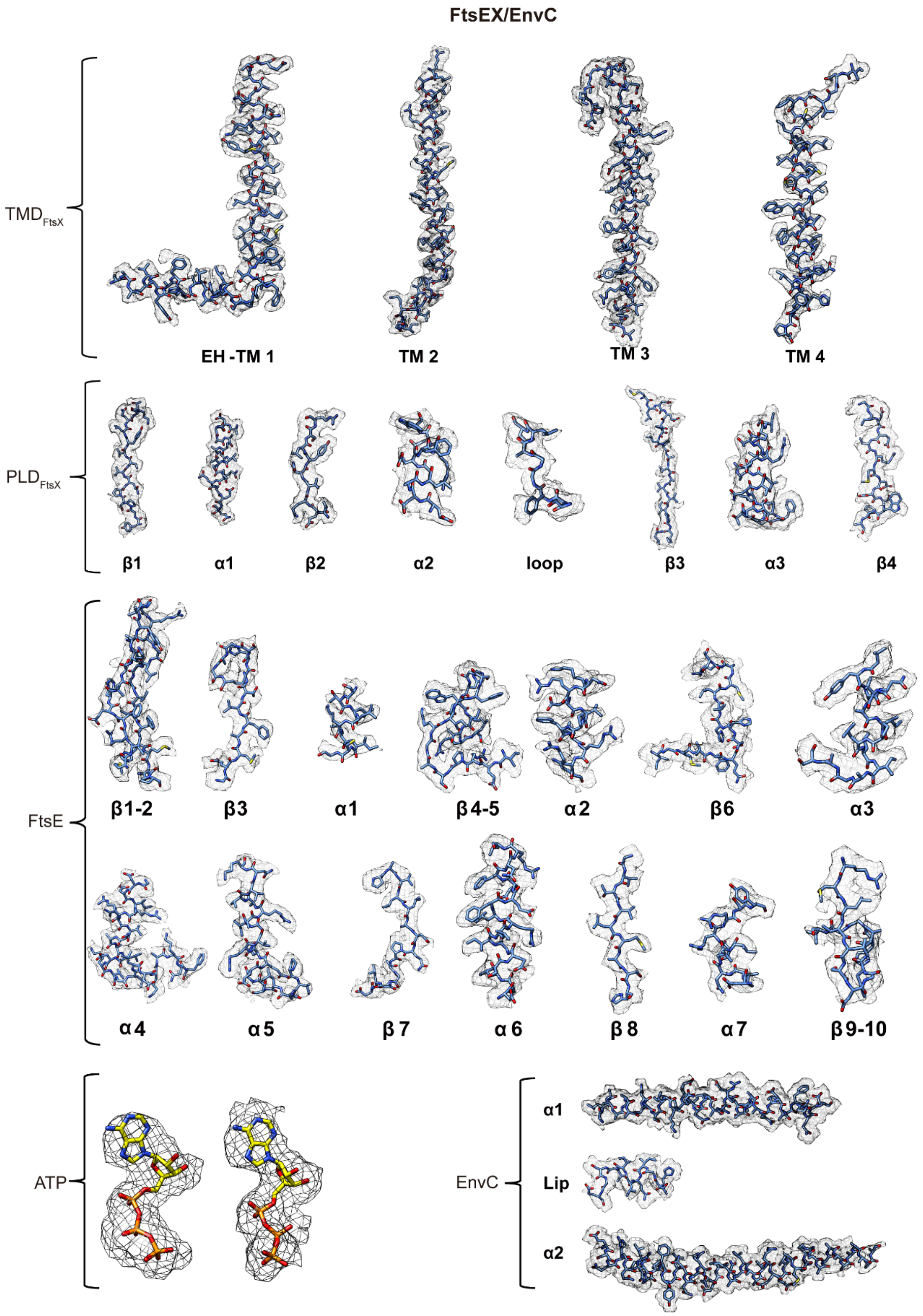


**Figure supplement 6. Cryo-EM density of different regions of FtsEX and EnvC in the structure of FtsEX-EnvC complex.**

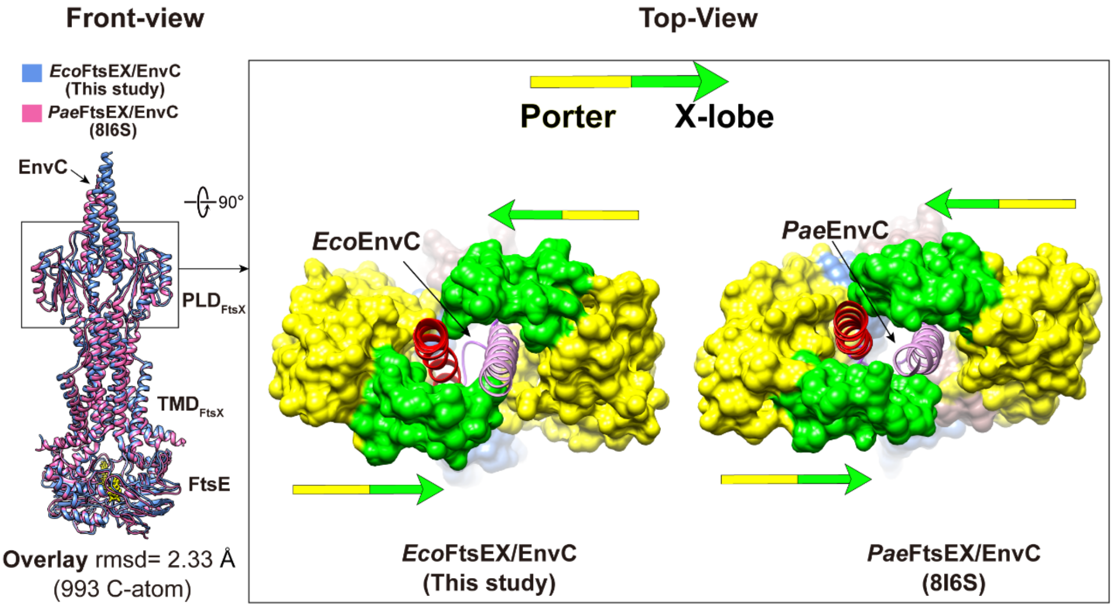


**Figure supplement 7. Structural comparison of PLD domain between *Eco*FtsEX/EnvC and *Pae*FtsEX/envC.** Left: Overlay of *Eco*FtsEX/EnvC and *Pae*FtsEX/envC (PDB: 8I6S) (1). *Eco*FtsEX/EnvC in blue, *Pae*FtsEX/envC in pink. Right: Surface representations of the PLD domains of *Eco*FtsEX/EnvC and *Pae*FtsEX/envC with top-view, ribbon representations of the EnvC. the α1 of EnvC in plum, α2 of EnvC in red, the Porter of PLD in yellow, and the X-lobe of PLD in green.

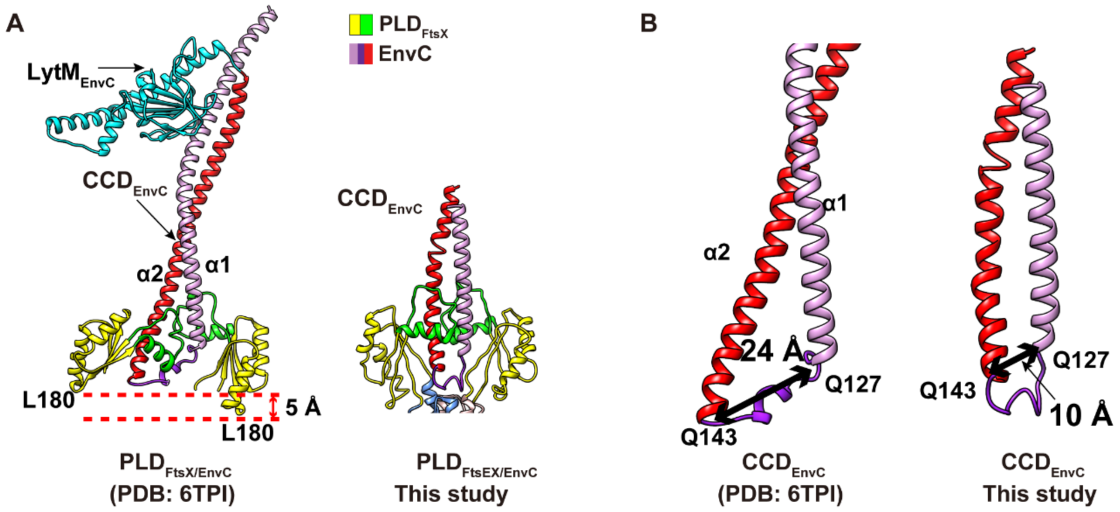


**Figure supplement 8. Structural comparison of cryoEM structure of FtsEX/EnvC complex and X-ray structure of PLD Domain with bound EnvC.** (**A**) Structural comparison highlighting the PLD/EnvC domains in both the FtsEX/EnvC complex and the PLD/EnvC structure (PDB: 6TPI) (2). (**B**) Structural comparison focusing on the CCD domain of EnvC between the FtsEX/EnvC complex and the PLD/EnvC structure. The α1 helix is shown in plum, the lip region in purple, and the α2 helix in red. Notably, both the PLD domain of FtsEX and the CCD domain of EnvC exhibit a more symmetrical conformation in the FtsEX/EnvC structure.

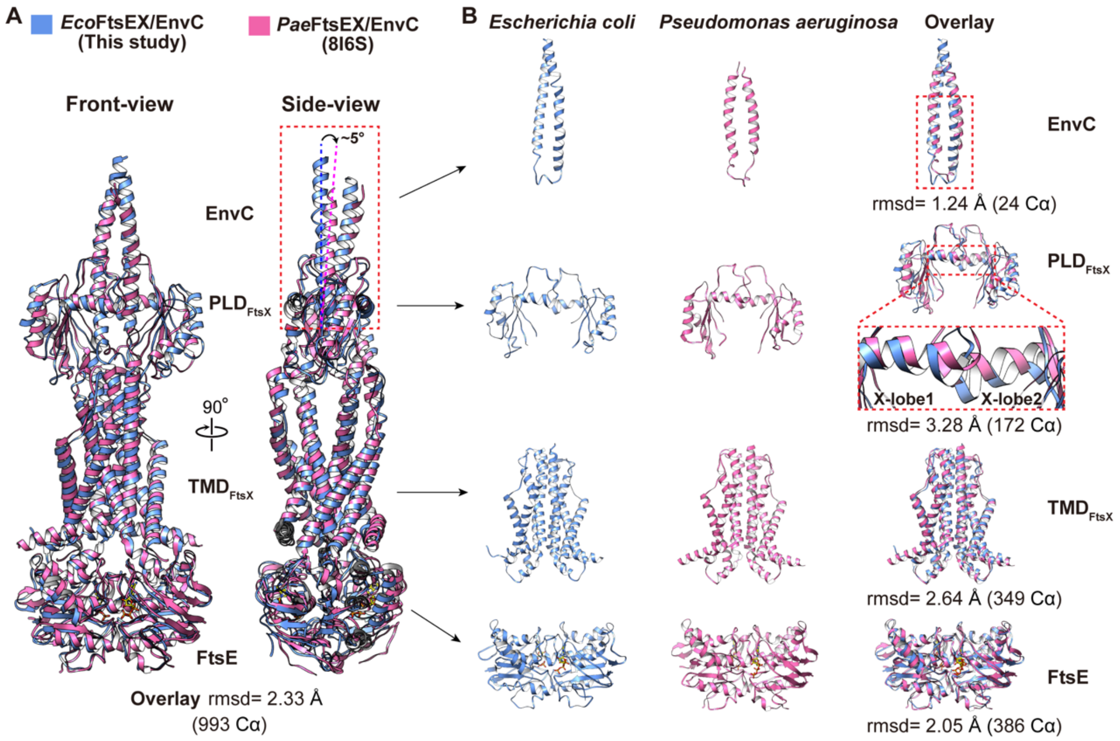


**Figure supplement 9. Structural comparison of *Eco*FtsEX/EnvC and *Pae*FtsEX/EnvC .** (**A**) Superposition of *Eco*FtsEX/EnvC and *Pae*FtsEX/EnvC (PDB: 8I6S) in front and side views, revealing a minor tilting difference in the bound EnvC. (**B**) Structural comparison of EnvC, the PLD and TMD regions of FtsX, and the NBD between *Eco*FtsEX/EnvC (blue) and *Pae*FtsEX/EnvC (pink). While the TMD and NBD regions are largely identical, differences are observed in the CCD of EnvC and the PLD domains. In *E. coli*, the two X-lobes of the PLD domain are positioned more tightly, pushing further toward the plasma membrane. This corresponds to a slightly more compact conformation between the two CCD coils in *E. coli* compared to *P. aeruginosa*.

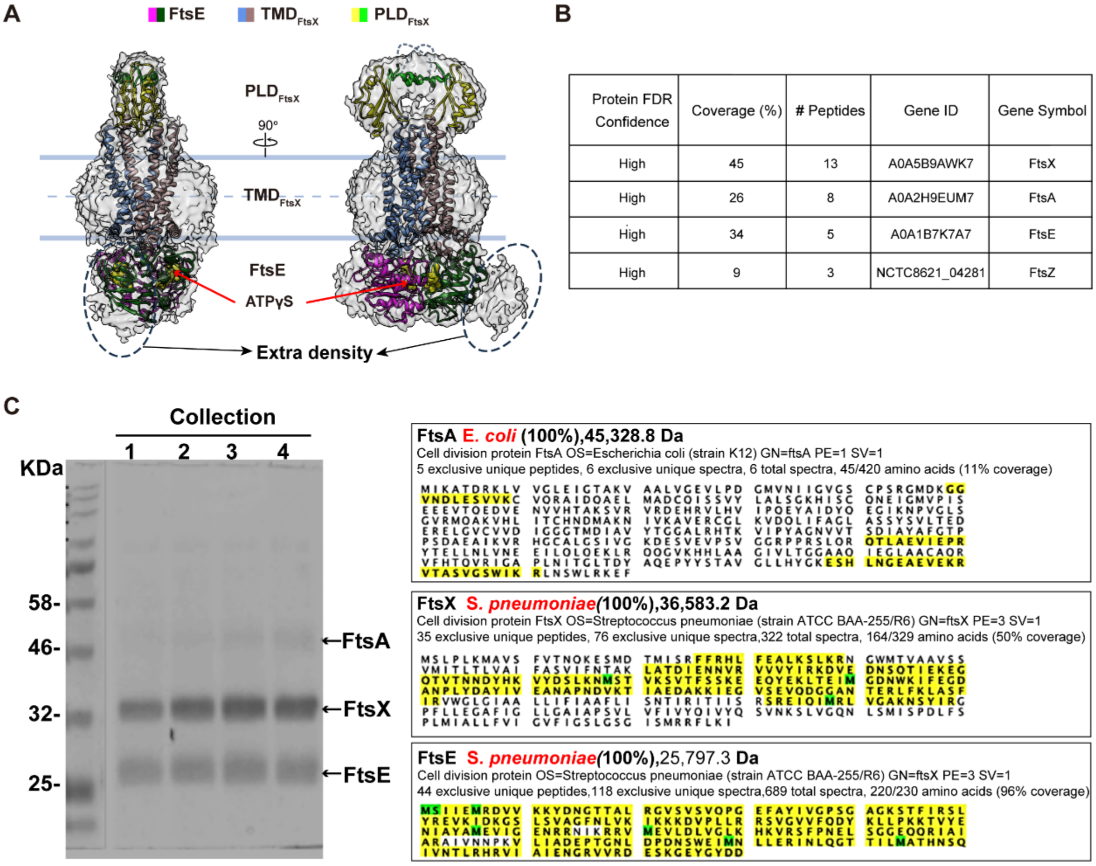


| Protein FDR Confidence | Coverage (%) | # Peptides | Gene ID | Gene Symbol |
|---|---|---|---|---|
| High | 45 | 13 | A0A5B9AWK7 | FtsX |
| High | 26 | 8 | A0A2H9EUM7 | FtsA |
| High | 34 | 5 | A0A1B7K7A7 | FtsE |
| High | 9 | 3 | NCTC8621_04281 | FtsZ |

**Figure supplement 10. Co-purification of endogenously FtsA/FtsZ with FtsEX from *E. coli*.** (**A**) Density map of FtsEX displayed at 70% transparency, shown in side- and front- views. Ribbon representation of FtsEX in the presence of ATPγS docked into EM density. Dotted lines outline the additional density. Color scheme: FtsE (magenta and dark green), FtsX (cornflower blue and rosy brown), Porter of $PLD_{FtsX}$ (yellow), and X-lobe of $PLD_{FtsX}$ (green). (**B**) Mass spectrometry results demonstrating the presence of considerable amounts of *Eco*FtsA and *Eco*FtsZ in the purified *Eco*FtsEX samples. (**C**) *Eco*FtsA copurified with *Sp*FtsEX. Left: SDS-PAGE analysis of the purified *Sp*FtsEX fractions. Right: Mass spectrometry analysis of SDS-PAGE bands from overexpressed *Sp*FtsE and *Sp*FtsX, along with endogenously copurified FtsA from *E. coli*. Yellow lines indicate sequence coverage by exclusive unique peptides. Coverage percentages of 6%, 44%, and 95% represent *Eco*FtsA, *Sp*FtsX, and *Sp*FtsE exclusive unique peptide coverage, respectively.

**Table supplement 1. Oligonucleotides used in this study.**

| Primer | Sequence (5'-3') | Template |
|---|---|---|
| **For construction of His$_6$- FtsEX** | | |
| *Eco*EX-WT F | accaccaccatatgattcgctttgaacat | Genomic DNA of *E. coli K12 Str MG1655* |
| *Eco*EX-WT R | tgctcgacaagcttttattcaggcgtaaag | |
| *Sp*EX-WT F | caccaccaccatatgtcaattattgaaat | Genomic DNA of *Streptococcus pneumoniae str. D39* |
| *SP*EX-WT R | ctcgacaagcttctaaatcttcaagaatcggc | |
| *Pae*EX-WT F | ccaccaccaccatatgatccgcttcgagcagg | Genomic DNA of *Pseudomonas aeruginosa ATCC 47085D* |
| *Pae*EX-WT R | gctcgacaagctttcaagtgatcccggcggcg | |
| *Bs*EX-WT F | ccaccaccatatgatagagatgaagga | Genomic DNA of *Bacillus_subtilis_subsp_subtilis_ATCC_23857*, |
| *Bs*EX-WT R | cgacaagcttttatactcgcagaaacttg | |
| **For construction of His$_6$-SUMO-EnvC** | | |
| EnvC-WT-F | tggtggatccgatgagcgtgaccaactc | Genomic DNA of *E. coli K12 Str MG1655* |
| EnvC-WT-R | tgctcgagtttatcttcccaaccacggct | |
| **For construction of His$_6$-SUMO-AmiB** | | |
| AmiB-WT-F | ggtggatccgcgacgctctctgatattc | Genomic DNA of *E. coli K12 Str MG1655* |
| AmiB-WT-R | gtgctcgagttagtttggcagcgtgcga | |
| **For FtsE mutation** | | |
| *Eco*FtsE-E163Q-F | ggcggaccaaccgactggtaacctggacg | Vector-*Eco*FtsEX-WT |
| *Eco*FtsE-E163Q-R | cagtcggttggtccgccagcagtaccgcg | |

**Table supplement 2. Plasmids used in this study.**

| **Plasmids** | **Promoter** | **Repressor** | **Fusion tags** | **Genes for overexpressed** | **Antibiotic resistance** | **Source** |
|---|---|---|---|---|---|---|
| pTD68 | T7 | lac repressor | His6 | *EcoFtsEX*, *EcoFtsE$^{E163Q}$X* *PaeFtsEX* *SpFtsEX* *BsFtsEX* | Ampicillin | (3) |
| pET28-Sumo | T7 | lac repressor | His6-Sumo | *EnvC, AmiB* | Kanamycin | (4) |

**Table supplement 3. Statistics of the cryo-EM structures presented in this study.**

| Cryo-EM data collection and processing | FtsEX/ATPγS | $FtsE^{E163Q}X$/EnvC/ATP |
|---|---|---|
| Voltage(kV) | 300 | 300 |
| Electron dose ($e^{-}/Å^{2}$) | 35 | 39 |
| Physical pixel (Å) | 1.06 | 1.06 |
| Number of movies | 3084 | 6054 |
| Number of particles for final map | 116,651 | 182,732 |
| Resolution (Å) | 3.7 | 3.4 |
| Map B-factor ($Å^{2}$) | -137 | -129 |
| **Model refinement** | | |
| Number of protein residues | 1021 | 1127 |
| Number of chains | 4 | 5 |
| Number of atoms | 7752 | 8555 |
| Ligands | 2 ATPγS | 2 ATP |
| **Geometric parameters (r.m.s.d.）** | | |
| Bond length (Å) | 0.004 | 0.005 |
| Bond angle (º) | 1.096 | 1.178 |
| **Ramachandran statistics** | | |
| Residues favoured (%) | 94.65 | 91.94 |
| Residues allowed (%) | 5.05 | 7.61 |
| Residues disallowed (%) | 0.30 | 0.45 |
| Rotamer outliers (%) | 0.13 | 0.12 |
| **MolProbity Score** | 1.69 | 1.79 |
